# Dissecting durum wheat time to anthesis into physiological traits using a QTL-based model

**DOI:** 10.1101/2023.02.25.530018

**Authors:** Pierre Martre, Rosella Motzo, Anna Maria Mastrangelo, Daniela Marone, Pasquale De Vita, Francesco Giunta

**Author notes:** Correspondence (PM).

## Abstract

Fine tuning crop development is a major breeding avenue to increase crop yield and for adaptation to climate change. In this study, we used a model that integrates our current understanding of the physiology of wheat phenology to predict the development and anthesis date of a RILs population of durum wheat with genotypic parameters controlling vernalization requirement, photoperiod sensitivity, and earliness *per se* estimated using leaf stage, final leaf number, anthesis date data from a pot experiment with vernalized and nonvernalized treatments combined with short- and long-day length. Predictions of final leaf number and anthesis date of the QTL-based model was evaluated for the whole population of RILs in a set of independent field trials and for the two parents, which were not used to estimate the parameter values. Our novel approach reduces the number of environments, experimental costs, and the time required to obtain the required data sets to develop a QTL-based prediction of model parameters. Moreover, the use of a physiologically based model of phenology gives new insight into genotype-phenology relations for wheat. We discuss the approach we used to estimate the parameters of the model and their association with QTL and major phenology genes that collocate at QTL.

**Highlight:** We used a modeling framework integrating our current understanding of the physiology of wheat phenology to dissect durum wheat time to anthesis into physiological traits and link them to QTL.

## Introduction

The increase in the occurrence and intensity of drought and heat stress due to global climate change is accompanied by a greater impact of genotype by environment interactions (G x E) on crop yields (Xiong *et al*., 2021), making breeding for adaptation more difficult. A fine-tuning of plant development is an avenue to cope with future climates and weather variability. Plant development is an important determinant of G x E and climate adaptation (Asseng *et al*., 2019; Fischer, 2016; Parent *et al*., 2018) and large and well understood genetic variations in vernalization, photoperiod sensitivity, and earliness *per se*, the three components of crop earliness, is available to crop breeders (Hyles *et al*., 2020; Kiss *et al*., 2017).

Ecophysiological models are powerful tools to get a better insight into how G x E interactions come about and to predict the performance of genotypes in defined environments (e.g. Bertin *et al*., 2010), although it requires more robust and biological sound crop models than do conventional agricultural applications (Hammer *et al*., 2019). Phenology models can be classified in two groups according to how they simulate development. The classical approach is based on accumulated thermal time between development phases modified by photoperiod and/or vernalization status of the plants. Alternately, a physiological approach dissects time to anthesis into primordium, leaf production, and leaf growth processes, which integrate the effects of vernalization and photoperiod (He *et al*., 2012; Jamieson *et al*., 1998). These two approaches can give similar predictions of anthesis date (Jamieson *et al*., 2007). However, the advantage of a physiological-based approach to dissect flowering time into component traits goes beyond the capability to simulate anthesis date by establishing a strong physiological link between phenotype and genotype (Brown *et al*., 2013).

The structure of a model and the way interactions between the underlying processes are considered is essential to model genetic variability (Parent and Tardieu, 2014). To correctly simulate G x E, model architecture and associated coefficients should capture and integrate the physiological basis of the genetic variations. The physiological-based approach to model plant development has a greater potential explanatory capability of G x E because it simulates the avenues by which each genotype reaches anthesis. Whether the same anthesis date is reached by two different genotypes through less leaves or through a faster rate of leaf appearance is likely to affect genotype adaptation, not only through time to anthesis, but also via processes like leaf growth and final leaf size (Dornbusch *et al*., 2011), tiller production and mortality (Giunta *et al*., 2018) or ear fertility (Gonzalez-Navarro *et al*., 2016; Ochagavía *et al*., 2018; Ochagavía *et al*., 2017). The physiological approach to model phenology allows linking phenology with leaf area and tillering and to analyze interactions and trade-offs between these processes (Abichou *et al*., 2018; Martre and Dambreville, 2018).

Previous studies linked crop phenology model parameters with known phenology genes (Hoogenboom and White, 2003; Hoogenboom *et al*., 1997, for common bean; White *et al*., 2008, for winter wheat; Zheng *et al*., 2013, for spring wheat) or by identifying quantitative trait loci (QTL) associated with model parameters (Bogard *et al*., 2020a, for spring wheat; Bogard *et al*., 2014, for winter wheat; Nakagawa *et al*., 2005, for rice; Yin *et al*., 2005, for spring barley). All these studies have used phenology models based on accumulated thermal time between growth phases that do not consider leaf development. ‘Genetic’ parameters of the models were estimated together using observations of heading or anthesis date, which imply a long phenotypic distance between the observed variables and the model parameters.

In this study we developed a QTL-based model based on the phenological framework proposed by Jamieson *et al*. (1998) to predict leaf development and anthesis date of a recombinant inbreed lines (RILs) population of durum wheat (*Triticum turgidum* L. subsp. *durum* (Desf.) Husn.). In contrast with previous studies, we estimated the parameters controlling vernalization requirement, photoperiod sensitivity, and earliness *per se* for each genotype separately using leaf stage, final number, anthesis date data from a pot experiment with vernalized and nonvernalized treatments combined with short- and long-day length. QTL associated with each of the five genetic parameters of the model were used to obtain multiple linear regression prediction of the parameter values. Predictions of final leaf number and anthesis date of the QTL-based model was evaluated for the whole population of RILs in a set of independent field trials and for the two parents, which were not used to estimate the parameter values. Our approach reduces the number of environments, experimental costs, and the time required to obtain the required data sets to develop a QTL-based prediction of model parameters. The use of a physiologically based model of phenology gives new insight into genotype-phenology relations for wheat. Several of the QTL associated with model parameters co-localized with known vernalization requirement and photoperiod genes or QTL.

## Materials and methods

### Plant materials

Ninety-one lines of a F2-derived, F8-F9 recombinant inbred lines (RILs) mapping population obtained from a cross between the Italian durum wheat (*Triticum turgidum* L. subsp. *durum* (Desf.) Husn.) cultivars Ofanto and Cappelli was used (Verlotta *et al*., 2010). Ofanto is an early flowering, semi-dwarf cultivar released in 1990 that originated from a cross between the durum wheat cultivars Appulo and Adamello. Cappelli is late flowering with vernalization requirement and tall cultivar released in Italy in 1915 derived the North-African landrace ‘Jean Retifah’. The two parents of the mapping population were also used in this study.

### Experimental treatments and phenotypic data used for parameter estimation

A pot experiment with a set of three treatments (LDV, long days vernalized; LDNV, long days nonvernalized; and SDV, short days vernalized) was conducted at Ottava, Sardinia, Italy (41° N 8° E; 225 m above sea level; Giunta *et al*., 2018; Sanna *et al*., 2014) to estimate the genetic parameters of the model. Seeds of similar size were imbibed for 24 h at room temperature on water saturated Whatman paper discs in Petri dishes. For the nonvernalized treatment, germinated seeds were directly transplanted in 5 L pots (three seeds per pot) filled with 1:2 (v:v) mixture of sand and sandy-clay-loam soil. For the two vernalized treatments, germinated seeds were transferred in a controlled-temperature cabinet where they were maintained for 40 days at 4°C in the dark. At the end of the vernalization treatments their coleoptile was about 3-cm long and the first seminal root was about 4-cm long. The two long day treatments were potted on 24 May and the short-day vernalized treatment was potted on 23 December of the same year. Two pots were used for each RIL/treatment combination and were arranged in a completely randomized design. The May-sown plants were maintained outdoors, and the December-sown ones were kept in a greenhouse. The pots were watered and fertilized as required. Daily weather data were recorded in a meteorological station located 300 m from the field, temperatures were recorded inside the greenhouse near the plants. The environmental conditions for the three treatments are summarized in Supplementary Table S1.

The plants were monitored twice weekly to record the number and length of the leaves which had appeared on the main stem, the appearance of the flag leaf ligule, and anthesis on main stem. Anthesis was recorded when 50% of the anthers on the ear of the main stem were visible (that is, Zadoks growth stage 69; Zadoks *et al*., 1974). The Haun stage (decimal leaf stage) was calculated following Haun (1973):

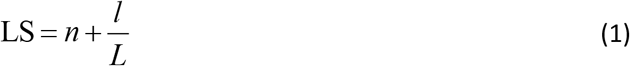

where *n* is the number of ligulated leaves, *l* is the exposed length of leaf *n*+1 at the time of measurement, and *L* is the final length of the blade of leaf *n*+1. The exposed length of a leaf was measured with a ruler as the distance from leaf tip to the upper collar of the sheath tube. Best linear unbiased predictors (BLUPs) were calculated for each RIL and trait from a mixed-model ANOVA as described in Sanna *et al*. (2014).

### Description of the wheat phenology model *SiriusQuality*

We used a modified version of the wheat phenology model described by He *et al*. (2012). The model is based on the framework proposed by Jamieson *et al*. (1998). It considers that vegetative and reproductive development is not independent and is coordinated and overlap in time (Kirby, 1990; Hay and Kirby, 1991). The successive appearance of leaves on the main-stem and tillers is the expression of the vegetative development, while anthesis is a particular stage in the reproductive development of the plant. Within this framework, the variations associated with vernalization requirement and daylength sensitivity are described in terms of primordium initiation, leaf production, and final main stem leaf number.

The leaf production phase is modeled based on two independently controlled processes, leaf initiation (primordia formation) and emergence (leaf tip appearance). The interaction between these processes leads to the determination of the final number of leaves (*Lf*) produced on the main stem. At any time during vegetative development the number of apex primordia (PN) is calculated through a metric relationship with leaf number under the assumption that the apex contains four primordia at plant emergence (PNini) and that they accumulate at twice the rate of leaf emergence (PNslope; Brooking and Jamieson, 2002):

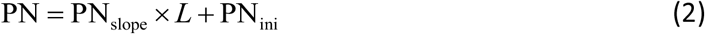

The rate of leaf appearance is described with a segmented linear model (Jamieson et al., 1995) where the first three leaves appear more rapidly than the next ones:

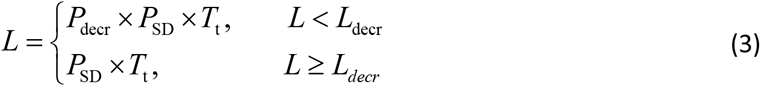

where *L* is the the number of appeared leaves on the main stem (equivalent to the Haun stage), *T*_t_ is the thermal time accumulated by the apex since plant emergence; *P*_SD_ is the phyllochron modified by sowing date for the first three leaves; *P*_decr_ is a factor (set at 0.75) decreasing the phyllochron for leaf number less than *L*_decr_; and *L*_decr_ is the Haun stage (set at 3 leaves) up to which *P* is decreased by *P*_decr_. Thermal time since plant emergence (*T_t_*) is calculated using a linear model of daily mean temperature with a base temperature of 0°C. Initially the controlling temperature (apex temperature) is assumed to be that of the near soil surface (0-2 cm), and then that of the canopy after Haun stage 4. Near soil surface temperature and canopy temperature are calculated using a surface energy balance model (Jamieson *et al*., 1995).

Many studies have shown that phyllochron depends on the sowing date (e.g. Baumont *et al*., 2019; McMaster *et al*., 2003; Slafer and Rawson, 1997). In *SiriusQuality*, for a winter sowing (day of the year 1 to 90 for the Northern hemisphere) the phyllochron decreases linearly with the sowing date and is minimum until mid-July for the Northern hemisphere (day of the year 200; He *et al*., 2012):

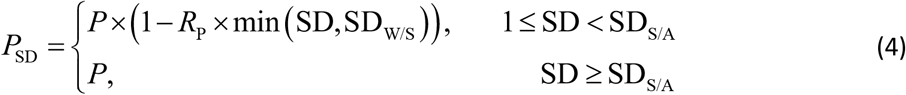

where SD is the sowing date in day of the year; *P* is the phyllochron for autumn sowing; *R*_P_ is the rate of decrease of *P*_SD_ for winter sowing; SD_W/S_ and SD_S/A_ are the sowing dates for which *P*_SD_ is minimum and maximum, respectively.

Vernalization progress and photoperiodic responses are modeled as sequential processes. Vernalization starts once the seed has imbibed water, which is assumed to take one day. In winter wheat, and other cereals, vernalization requirement can be eliminated or greatly reduced by a prolonged exposure to short daylength (Dubcovsky *et al*., 2006; Evans, 1987), a process referred as short day vernalization. We modified the vernalization model described by He *et al*. (2012) to account for this process. The photoperiodic effect on the vernalization rate is likely to involve a quantitative 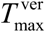 interaction with temperature rather than a complete replacement of the vernalization requirement (Brooking & Jamieson, 2002; Allard et al., 2012). In the revised model, the daily vernalization rate (*V*_rate_) increases at a constant rate (VAI) with daily mean temperature from its value (VBEE) at the minimum vernalizing temperature 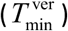 to a maximum for an optimum temperature 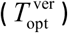. For temperature above 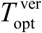, under short days, *V*_rate_ reduces to zero at the maximum vernalizing temperature 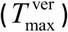, while under long days, *V*_rate_ stays at its maximum value. The effectiveness of short days decreases progressively as photoperiods increases. *V*_rate_ is given by:

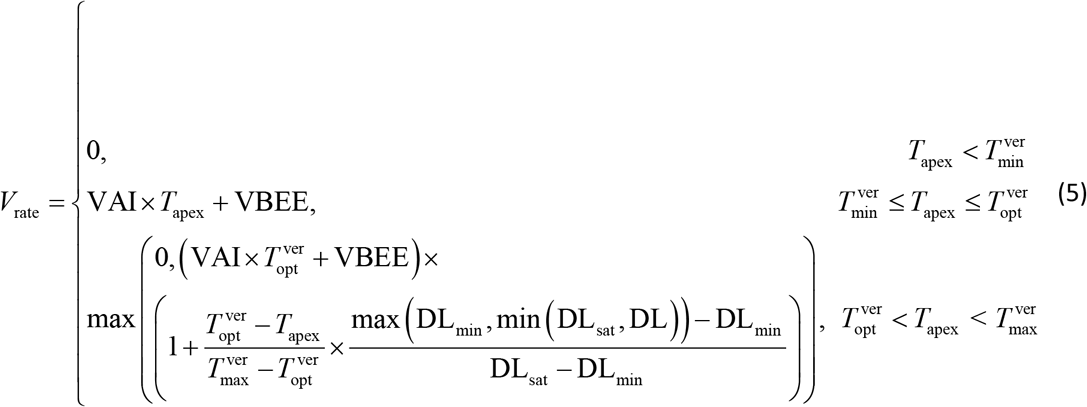

where *T*_apex_ is the apex temperature, DL is the day length of the current day, and DL_sat_ and DL_min_ are the saturation and minimum daylength for short day vernalization, respectively. The progress toward full vernalization (*V*_prog_) is simulated as a time integral:

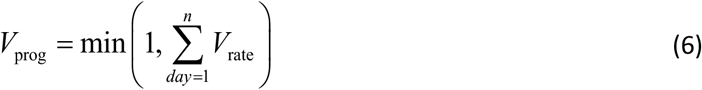

Two parameters define the minimum 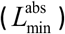 and maximum 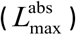 number of leaves that can be initiated on the main stem. The model assumes that plants start with a high potential leaf number (*L*_pot_ set to an initial value of 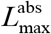) which decreases with vernalization progress:

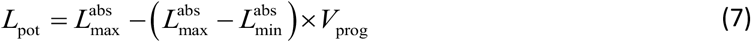

Vernalization is complete when one of the following three conditions is met: (1) *V*_prog_ equals 1; (2) *L*_pot_ equals 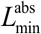; or (3) *L*_pot_ equals PN. All the primordium formed during the vernalization phase are assumed to produce leaves. 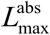 corresponds to the number of leaves produced by a winter genotype grown under long days at a temperature above 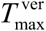.

The plant responds to DL only once vernalization is completed. Daylength sensitivity leads to an increase in the number of leaf primordia resulting from the vernalization routine. If DL of the day when vernalization is completed exceeds a given value (DLsat), the final leaf number on main stem (*L*_f_) is set to the value calculated at the end of the vernalization routine and the floral initiation is reached. For DL shorter than DLsat, Brooking *et al*. (1995) have shown that *L*_f_ is determined by DL at the stage of two leaves after the flag leaf primordium has been formed. This creates the need for an iterative calculation of an approximate final leaf number (*L*_app_) that stops when the required leaf stage is reached:

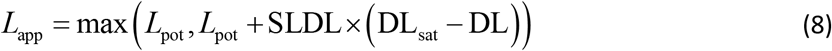

where, SLDL is a parameter defining the day length response as a linear function of DL. It is assumed that the attainment of the stage “two leaves after flag leaf primordium” is reached when half of the leaves have emerged (Brooking *et al*., 1995):

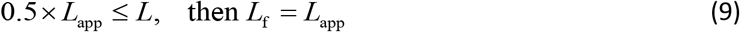

When this condition is fulfilled, transition to floral initiation is completed and *L*_f_ is equal to the number of primordia formed on that day. Anthesis occurs a fixed number of phyllochron (PFLLAnth) after the appearance of the flag ligule.

The model described above has been developed as an independent executable component (Manceau and Martre, 2018) in the BioMA software framework (Donatelli and Rizzoli, 2008) integrated in the wheat model *SiriusQuality*, version 2.0.57777 (He *et al*., 2012; Martre and Dambreville, 2018; Martre *et al*., 2006).

### Estimation of the ecophysiological model parameters

Five parameters of the phenology model were estimated for each of the 91 RILs using the three treatments of the pot experiment described above (Table 1). These parameters were estimated based a previous study which showed that *P*, SLDL and VAI are enough to predict genetic variability of winter wheat genotypes (He *et al*., 2012; Rincent *et al*., 2017). PFLLAnth and 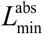 were also estimated because a previous analysis of the data set used for parameter estimation in this study revealed a significant genetic variability for these two traits (Sanna *et al*., 2014).

**Table 1.**
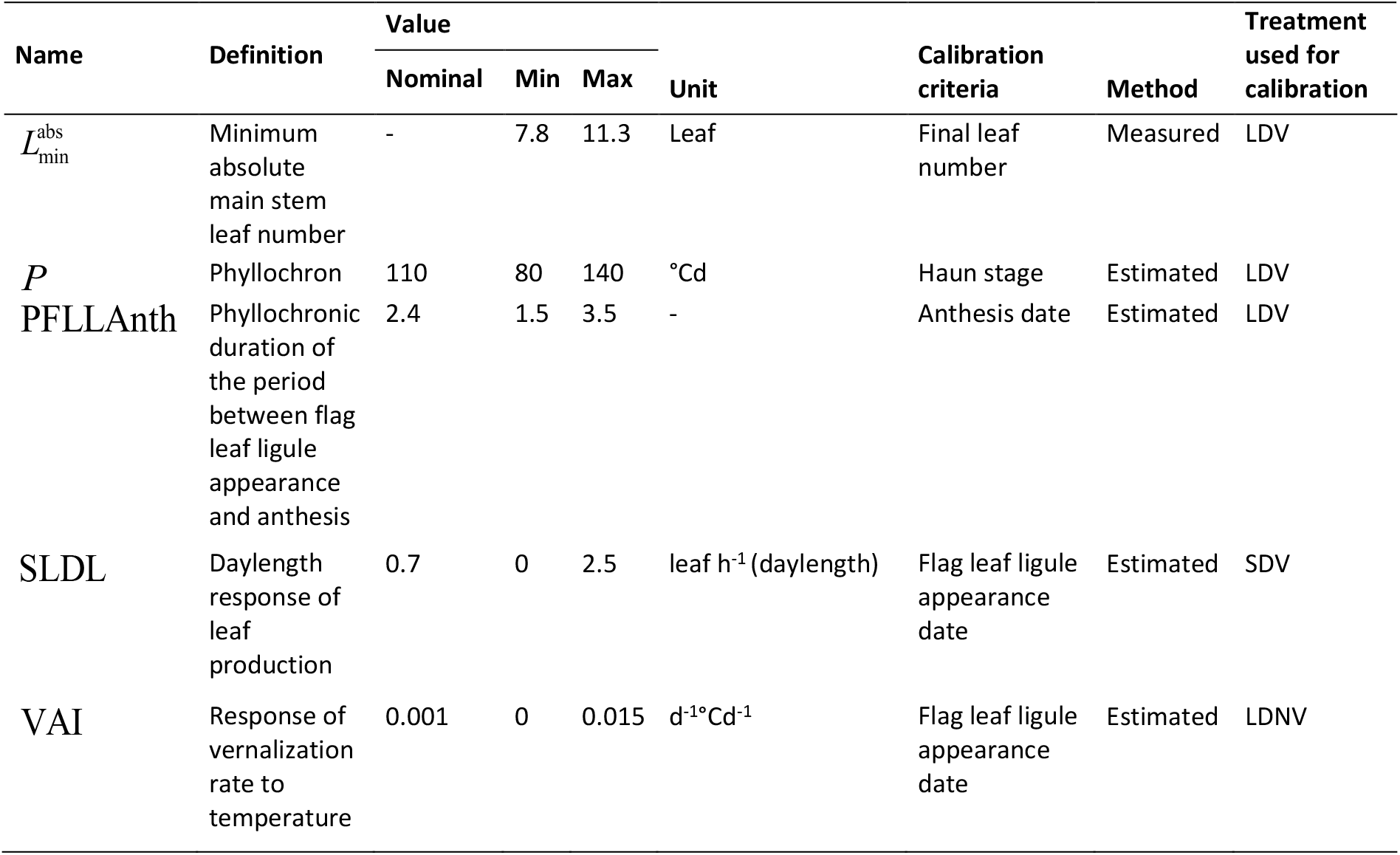
Name, symbol, definition, nominal, minimal, and maximal value, unit and calibration criteria of the calibrated genetic parameters of *SiriusQuality* phenology sub-model. The four parameters were optimized squentially in order they are shown in the table.

We designed a calibration procedure that minimizes the interactions between the different components of phenology. First, three parameters controlling earliness *per se* (*P*, 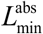, PFLLAnth) were estimated with the LDV treatment. 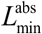 was set equal to the measured value of *L*_f_, then *P* and PFLLAnth were estimated sequentially by minimalizing the root mean squared error (RMSE) for Haun stage and the absolute error (AE) anthesis date, respectively. Then the sensitivity to daylength (SLDL) was estimated by minimizing the AE for the date of flag ligule appearance for SDV treatment. Finally, the slope of the response vernalization rate to temperature (VAI) was estimated by minimizing the AE for the date of flag ligule appearance for LDV treatment. Parameters were estimated with the Brent hybrid root-finding algorithm (Brent, 1973) by using the ‘optim’ function of the ‘stats’ package of the R software program, version 4.1.3 (R Core Team, 2022). The other parameters of the model were set to the values given by He *et al*. (2012), except *L*_decr_, 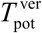 and 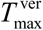 which were increased following the work of Brown *et al*. (2013) and VBEE that was also increased following Robertson *et al*. (1996) to take into account the lower response of vernalization rate to temperature for durum wheat compared with winter bread wheat(Supplementary Table S2). All simulations started on the sowing date.

### Genetic map and quantitative trait loci detection

An updated version of the Ofanto × Cappelli genetic map previously reported (Marone *et al*., 2012) was developed and used for QTL analysis of the parameter values. Whole-genome profiling was performed using the DArT-Seq™ technology (Diversity Arrays Technology Pty Ltd, Australia). DArT-Seq™ detects both single nucleotide polymorphisms (SNPs) and presence–absence sequence variants, collectively referred to as DArT-Seq™ markers. Briefly, the genetic map is composed of 32 linkage groups which cover all of the chromosomes except 1A. The total number of markers is 9,267, of which 4,033 on the A genome and 5,594 on the B genome. The number of markers per chromosome ranges from 162 (4B) to 1,217 (6B). The map length spanned 2,119.2 cM, with 965.5 cM for the A genome, and 1,153.7 cM for the B genome.

QTL analysis was performed using the Composite Interval Mapping method (Zeng, 1994) with the Qgene software, version 4.3.10 (Joehanes and Nelson, 2008). Scanning interval of 1 cM between markers and tentative QTL with a window size of 10 cM was used to detect QTL. Marker cofactors for background control were set by single marker regression and simple interval analysis with a maximum of five controlling markers. Major QTL were defined as two or more linked markers associated with a parameter with a logarithm of odds (LOD) score > 5.0 and a phenotypic variance contribution > 10%. QTL with a LOD score > 2.8 and a phenotypic variance contribution < 10% were defined as moderate QTL. Tentative QTL with a LOD score between 1.0 and 2.8 were also considered for the prediction of QTL-based parameters. For main QTL effects, the positive sign of the estimates indicates that Ofanto allele contributed to the higher values of the parameter. The intervals of the QTL and flanking markers were determined following the method described by Darvasi and Soller (1997). The proportion of phenotypic variance explained by a single QTL was determined by the square of the partial correlation coefficient (*r*^2^). Graphical representation of linkage groups was carried out using the MapChart software, version 2.2 (Voorrips, 2002).

The available sequences of DArT-seq markers (provided by Triticarte, www.diversityarrays.com) were used as queries in a BLAST against the ‘Svevo’ genome (Maccaferri *et al*., 2019) to assign a physical interval to QTL identified in the present study. Similarly, available sequences of known genes involved in flowering time control in wheat and other species were used as queries in a BLAST search to identify their physical position onto the ‘Svevo’ genome. Physical position on the ‘Svevo’ genome of common markers mapped in previously published studies was also used for comparison with known QTL for phenological traits in tetraploid wheat.

### Quantitative trait loci prediction of the phenology model parameters

QTL-based values for each of the five estimated parameters were estimated for each RIL considering only additive QTL actions. Our aim was to be built a predictive model, therefore, all QTL with LOD score > 1 were considered. Following the approach used by Bogard *et al*. (2014), linear models for the five calibrated ecophysiological parameters were obtained using multiple linear regressions with backward elimination of the QTL by fitting the following statistical model to the estimated parameters values:

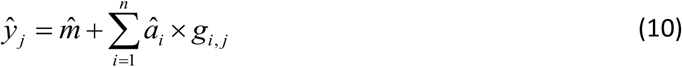

where 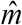 is the estimated intercept, *â_i_* is the estimated additive effect of the *i*-th QTL on the phenology model parameter, and *g_i,j_* is the allele of the *j*-th RIL at the *i*-th QTL. The Ofanto alleles were coded +1 and those of Cappelli −1.

### Field experiment for original and QTL-based model validation

Estimated and QTL-based values of the five parameters were used to simulate the development of the 91 RILs grown in the field during the 2012-2013 growing seasons at Ottava (experiment names OT13) and during the 2007-2008 (FO08) and 2008-2009 (FO09) growing seasons at Foggia, Italy (41.46° N, 15.55° E, 76 m a.s.l.). In Foggia, each line was planted at a rate of 40 seeds per row (1-m long) with 0. 3-m interrow spacing in a randomized complete block design with three replications. In Ottava, the RILs were sown with a 6-row planter at a density of 350 viable seeds m^-2^. Each plot consisted of six rows with an interrow spacing of 0.18 m and had a surface area of 10 m^2^. These three experiments were not used for parameter estimation. Anthesis dates was recorded at Ottava for each line and the two parents, while at Foggia heading date was recorded and anthesis date was estimated from the relationship obtained with OT13 data between thermal time to anthesis and thermal time to heading (*r*^2^ = 0.95, *P* < 0.001). Haun stage, final leaf number, flag leaf ligule appearance and anthesis dates were also recorded at Ottava using the protocol described above for the pot experiment. For FO08 and F09 means of anthesis were calculated, while for OT13 BLUPs were calculated for each RIL and trait from a mixed-model ANOVA as described in Sanna *et al*. (2014). Predictions using the QTL-based model parameters were compared with predictions using the estimated (original) parameters.

The QTL-based based model was also evaluated for the two parents, which were not used for QTL analysis, in the three environments described above, and in five (Cappelli) or 15 (Ofanto) other site/year/sowing date combinations. Cappelli was grown during the 2003-2004 growing season at Ottava with late-November and mid-February sowing dates and during the 2004-2005 growing season with early-January and mid-March sowing dates, and at Oristano, Sardinia, Italy (40° N, 8° W, 15 m a.s.l.) with mid-January sowing date. Ofanto was grown for eight consecutive years (harvests 1992 to 1999) at Ottava with sowing dates between mid-November and early-January, and at Oristano for seven years (harvests 1993 to 2000) with sowing dates between late-November and early-February. In all experiment, crops were sown at a density of 350 viable seeds m^-2^. Each plot was 7-m long with 8-rows and an interrow spacing of 0.18 m. The experimental design was a randomized complete block design with three replicates. The sowing dates and summary environmental conditions for all the trials are given in Supplementary Table S1. All trials were rainfed and other crop inputs including pest, weed and disease control, and nitrogen, potassium, and phosphate fertilizers were applied at levels to prevent nutrients or pests, weeds, and diseases from limiting plant development and growth. All crops were simulated from the day of sowing. At each site, daily weather data were recorded from meteorological stations located in the experimental farms near the experimental fields. For each parent, parameters values were obtained from the corresponding model linking genetic markers to model parameters and the model was used to predict the anthesis date.

### Statistics for model evaluation

Several statistics were calculated to assess the quality of the model simulation results. The observed and simulated data were compared using ordinary least square regression and the mean squared error (MSE). To get a better understanding of the model errors, the MSE was decomposed in non-unity slope (NU), squared bias (SB) and lack of correlation (LC) following Gauch *et al*. (2003). Spearman’s rank correlation coefficient was also calculated. All data analysis and graphs were done using R statistical software program version 4.2 (R Core Team, 2022).

## Results

### Estimations of the genetic parameters of the phenology model

The five estimated parameters showed large genetic variability between the RILs and significant transgressive segregation (Fig. 1). Ofanto and Cappelli had close values for *P* and SLDL. VAI was the most different parameter between the parents, with Cappelli having a much lower value than Ofanto. VAI had a clear bimodal distribution and the two parents had values close to the two peaks of the distribution. PFLLAnth was significantly correlated with *P* and SLDL (*r* = 0.40 and −0.27, respectively). The strongest correlation between parameters was between 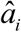 and SLDL (*r* = −0.66), although 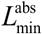 was measured in the LDV treatment and SLDL was estimated with the SDV treatments.

**Figure 1.**
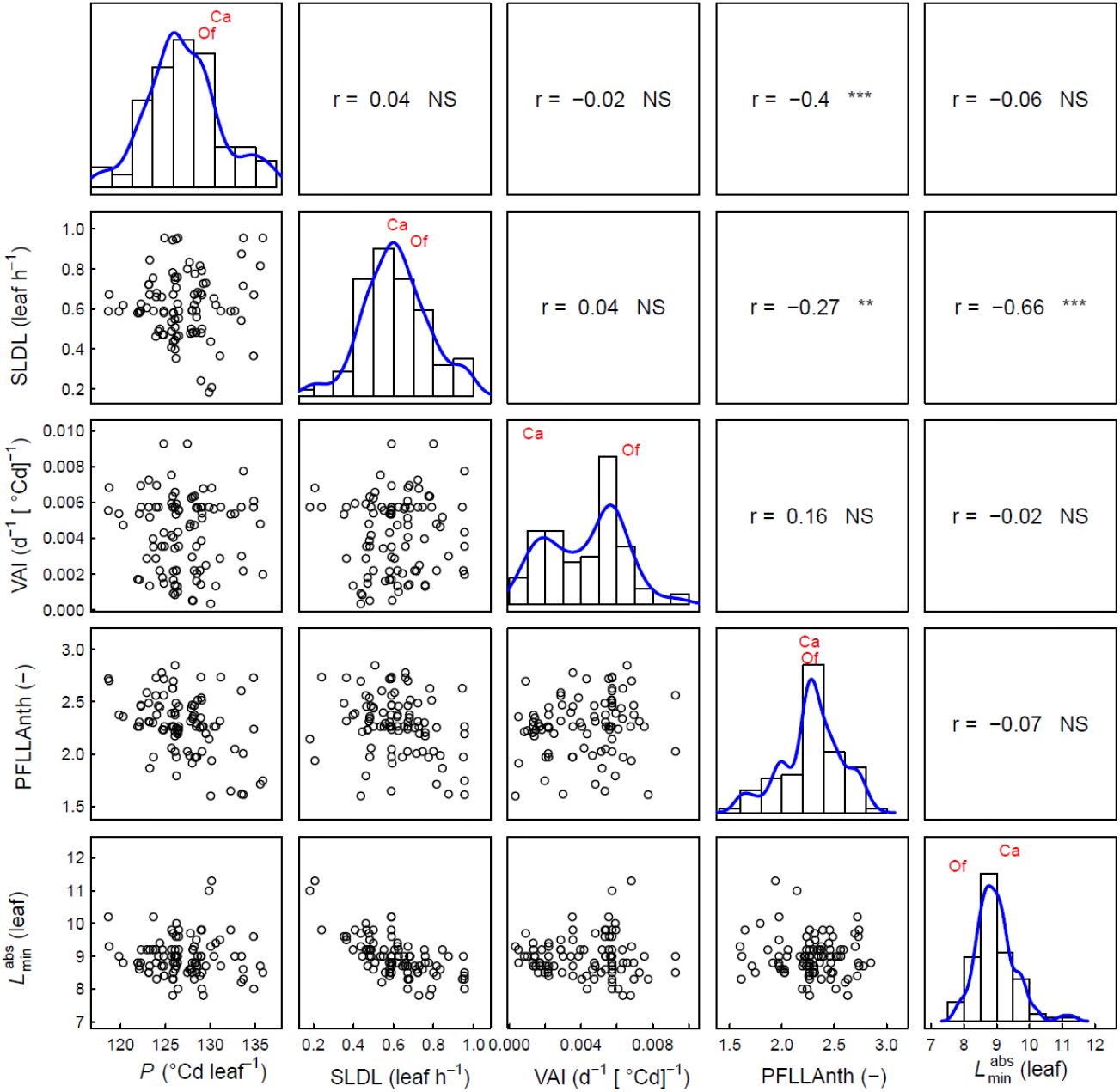
Distribution and correlations between the genetic parameters of *SiriusQuality* phenology model for 91 RILs of the Ofanto (Of) × Cappelli (Ca) cross. The phyllochron (*P*), the sensitivity to day length (SLDL), the response of the vernalization rate to temperature (VAI), and the number of phyllochron between flag leaf ligule appearance and anthesis (PFLLAnth) were estimated sequentially each using one of the three environments of the calibration dataset, while 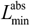 was measured in the LDV treatment. Correlation coefficients are reported above the diagonal. NS, not significant, ** *P* < 0.01, *** *P* < 0.001.

### Quantitative trait loci analysis and QTL-based prediction of model parameters

The genetic analysis of the estimated parameter values identified 13 moderate and major QTL (Table 2). All these QTL colocalized with known QTL for wheat phenology (Table 2). The percentage of variance of the parameters explained by each QTL varied between 14% (QTL 3 for *P*) and 44% (QTL 15 for VAI). No major or moderate QTL was identified for PFFLAnth but several tentative QTL colocalized with known QTL, including a QTL (QTL29, LOD = 2.0) previously identify for daylength sensitivity of heading date for winter wheat (Table 2). Two (for VAI) to five (for 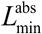) moderate or major QTL were identified for each of the other four parameters. Only one of these, QTL28, was associated with two model parameters (SLDL and 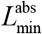), the other moderate and major QTL were associated with only one model parameter, but QTL2 (for 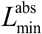) and QTL27 (for *P*) included a tentative region for SLDL (Fig. 3).

**Table 2.**
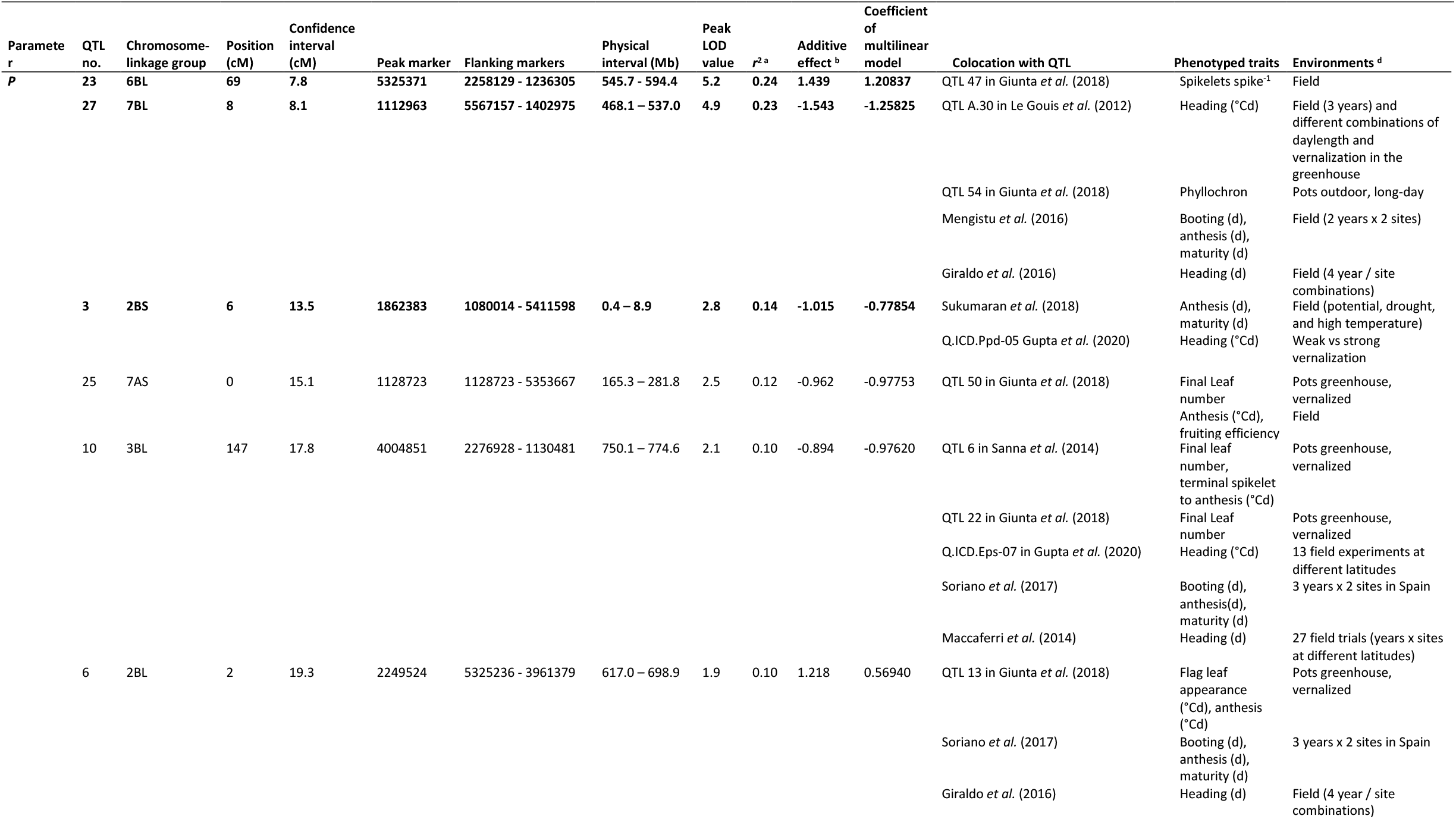
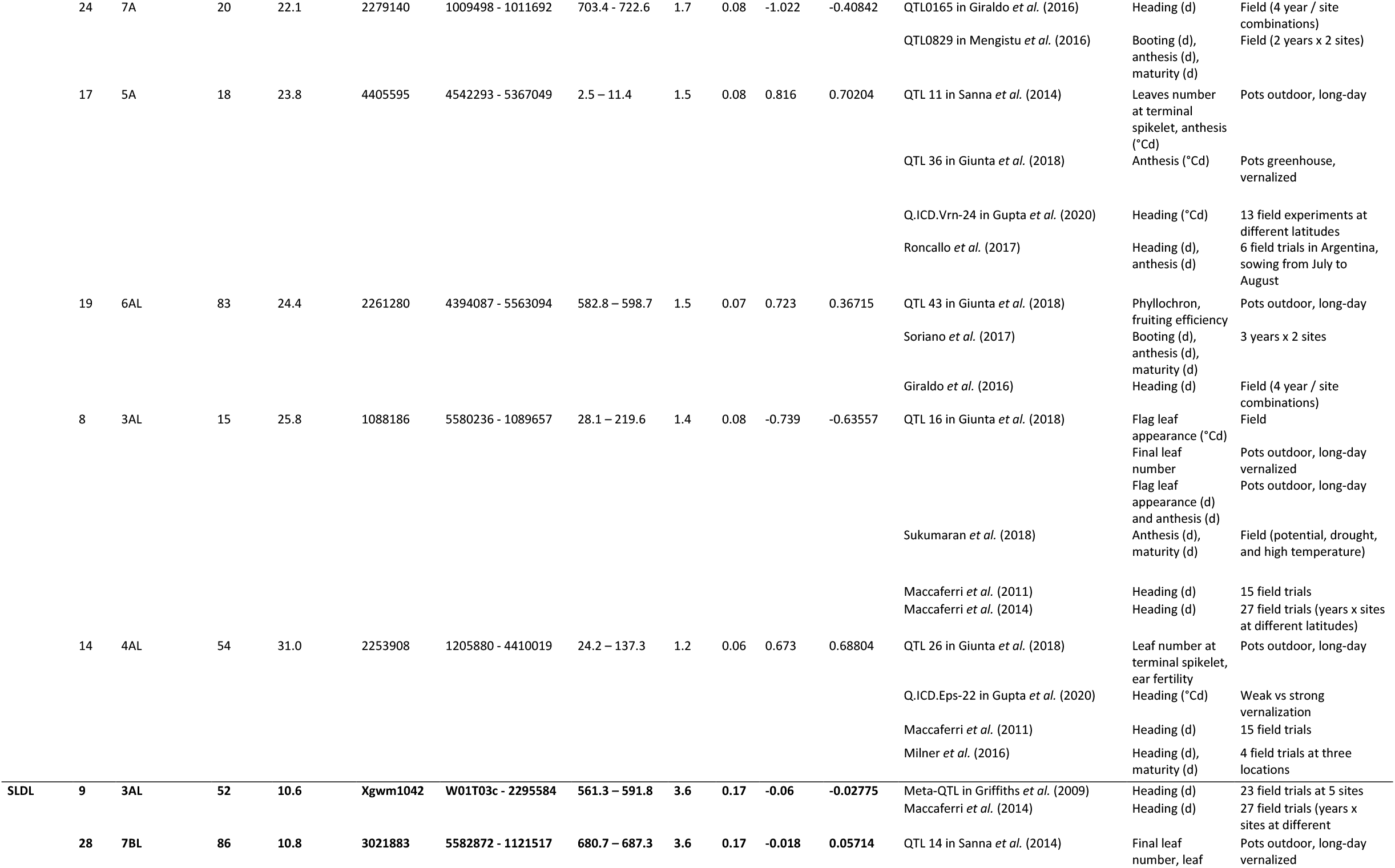
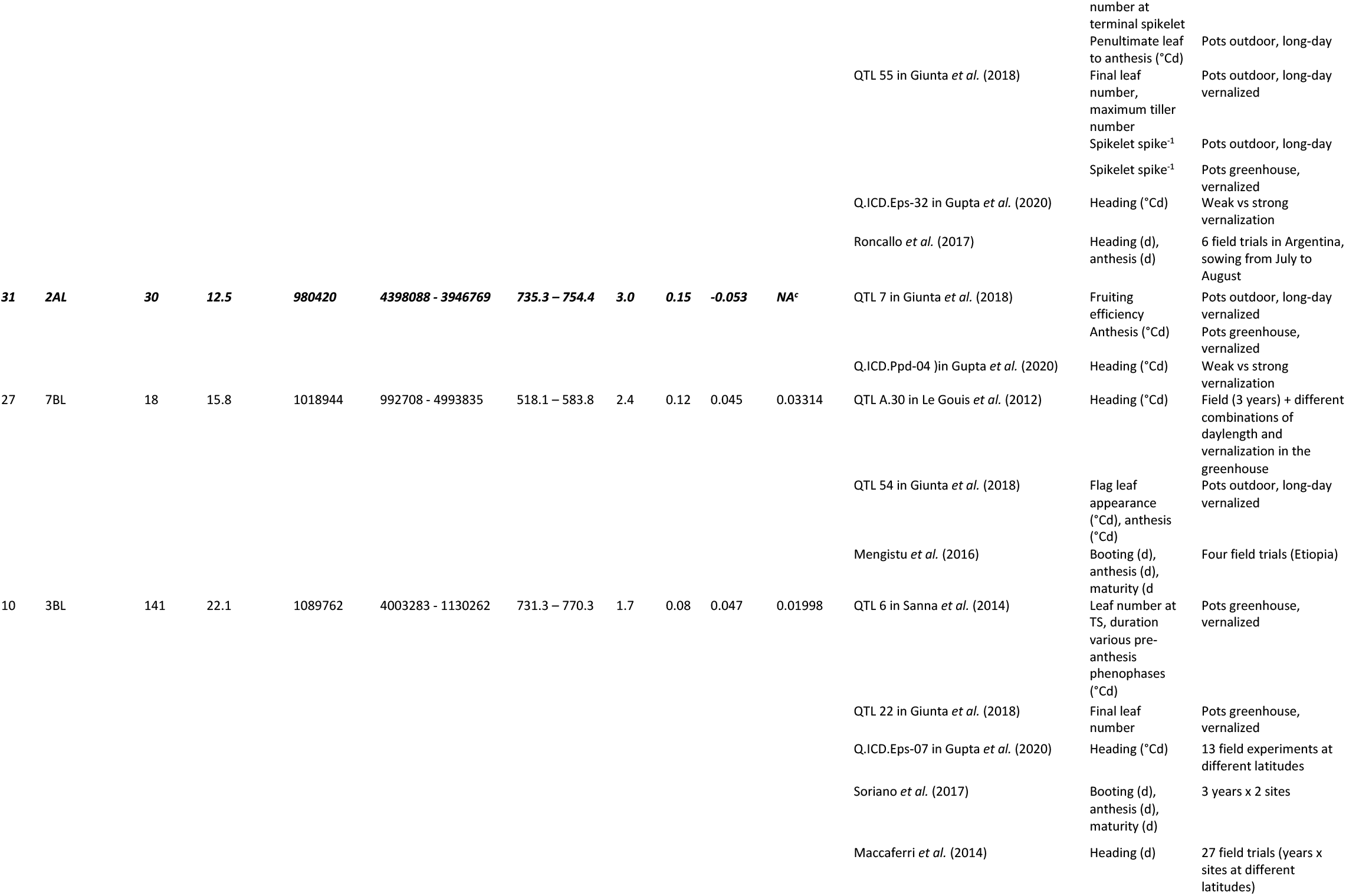
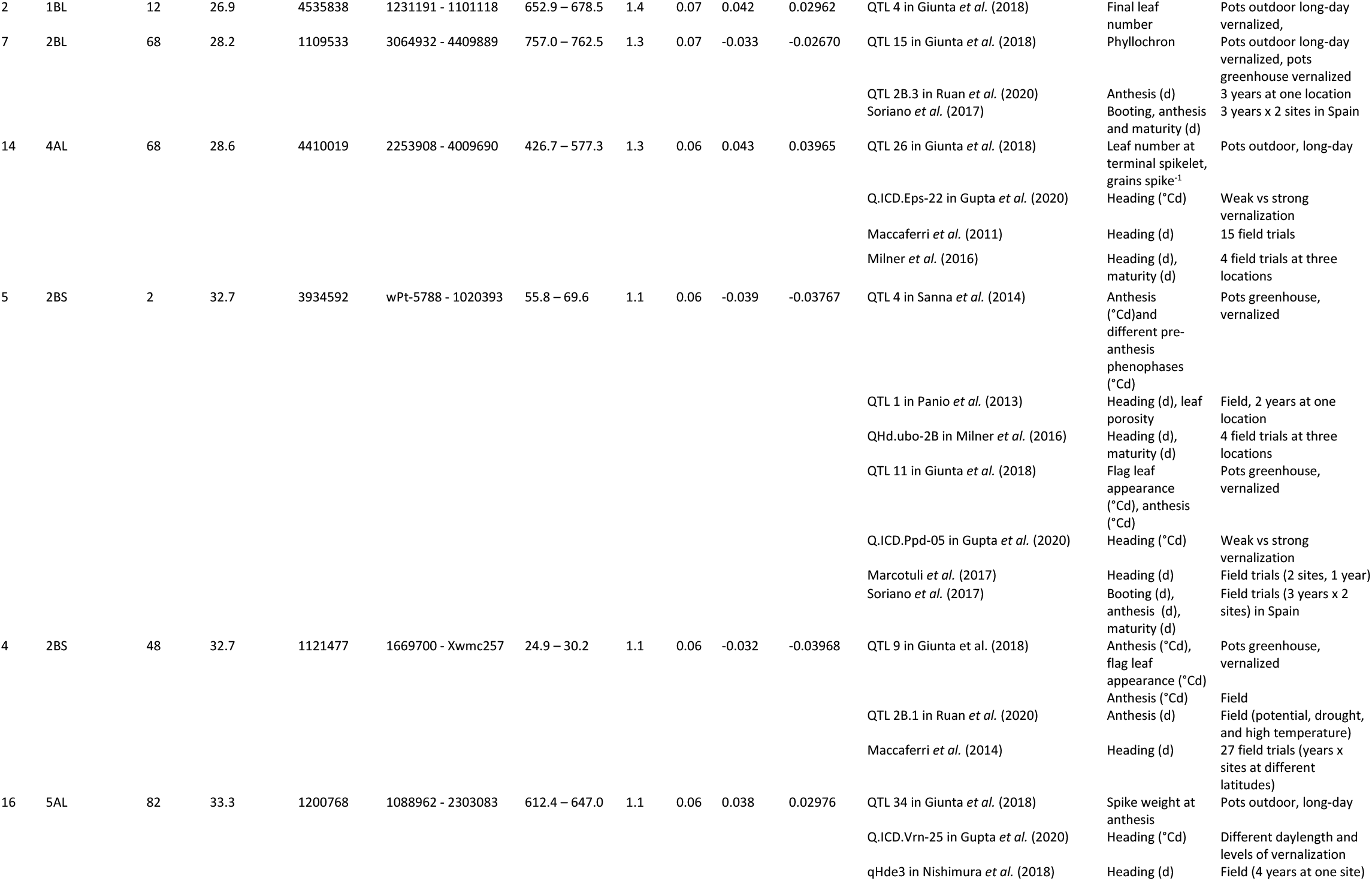
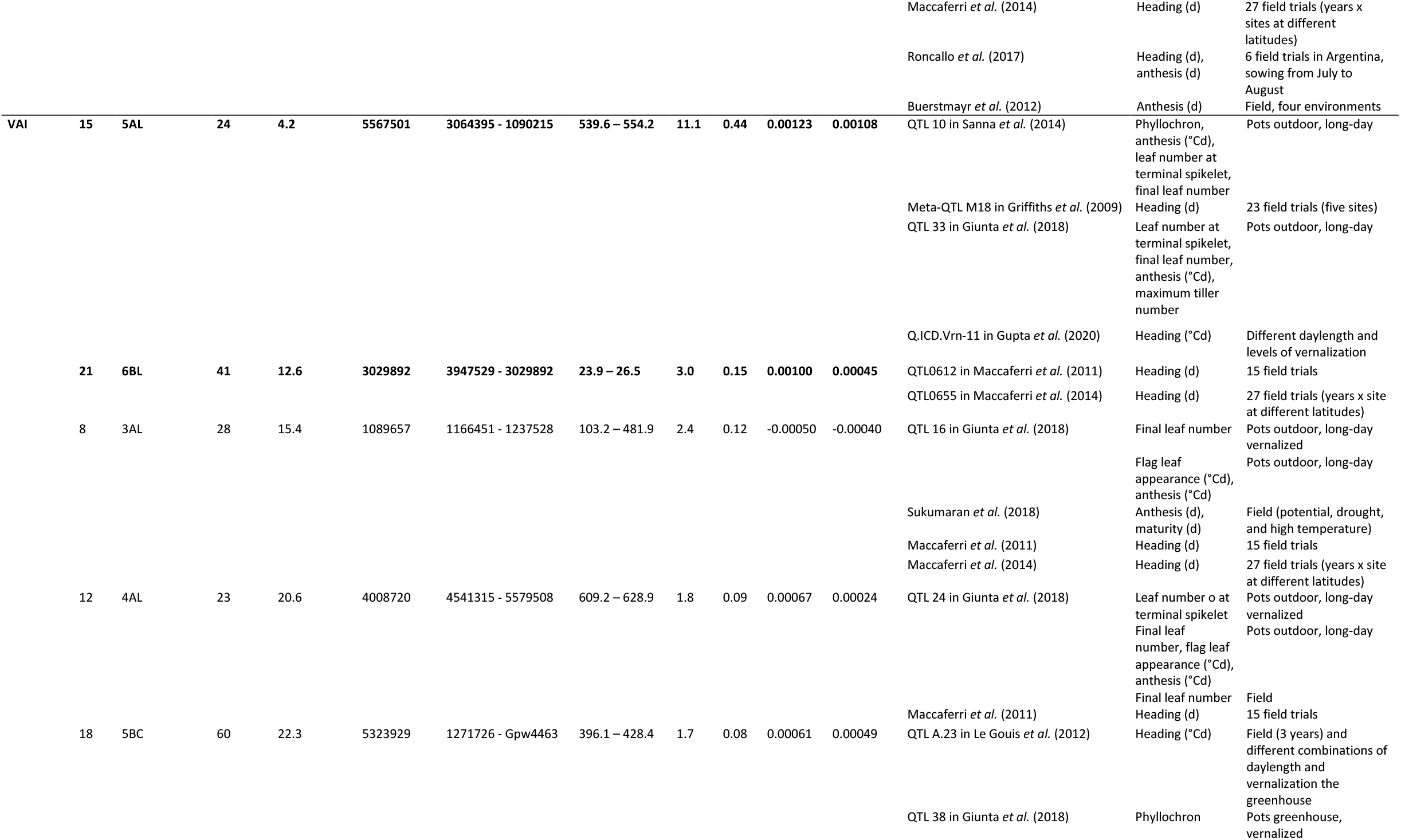
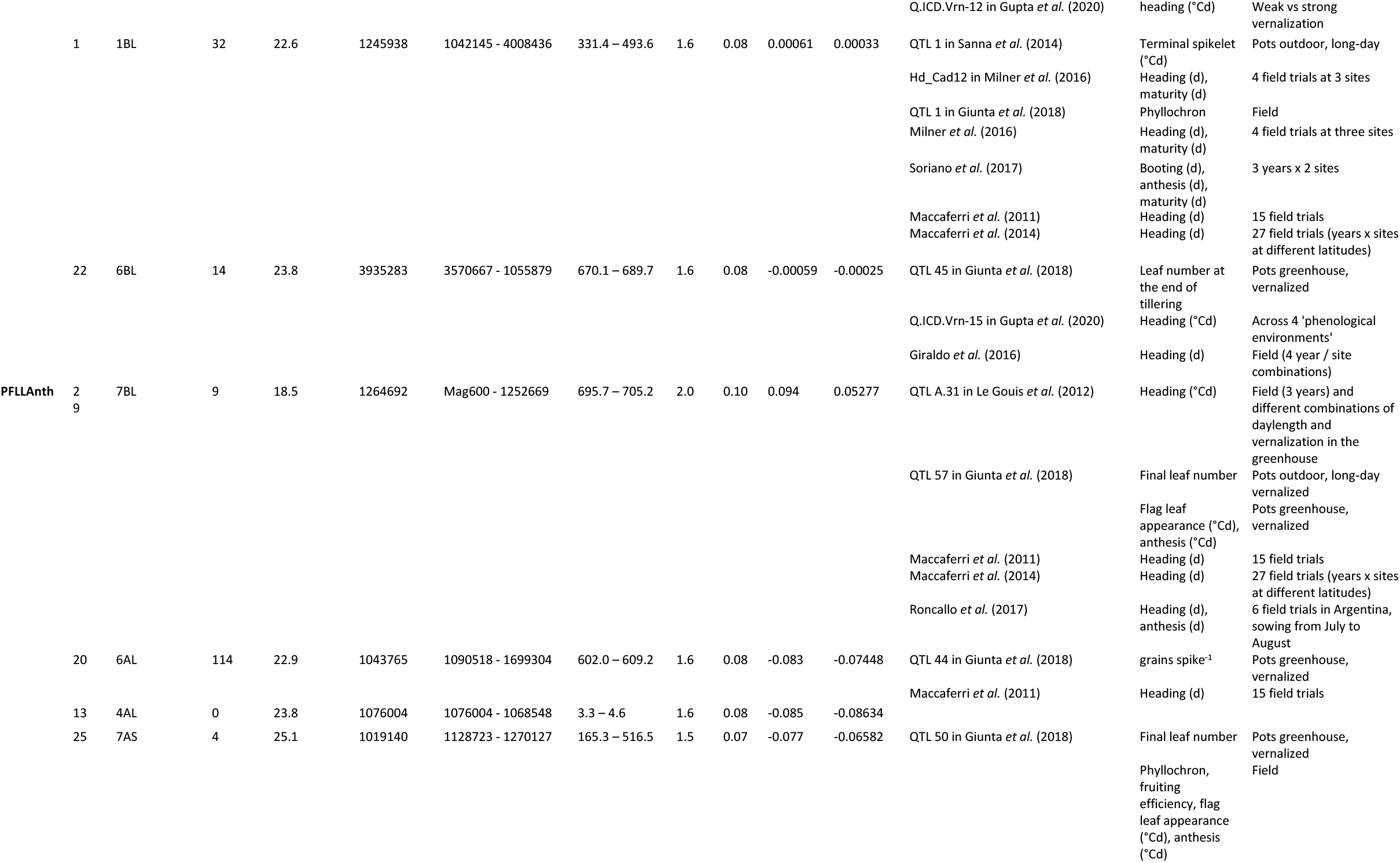
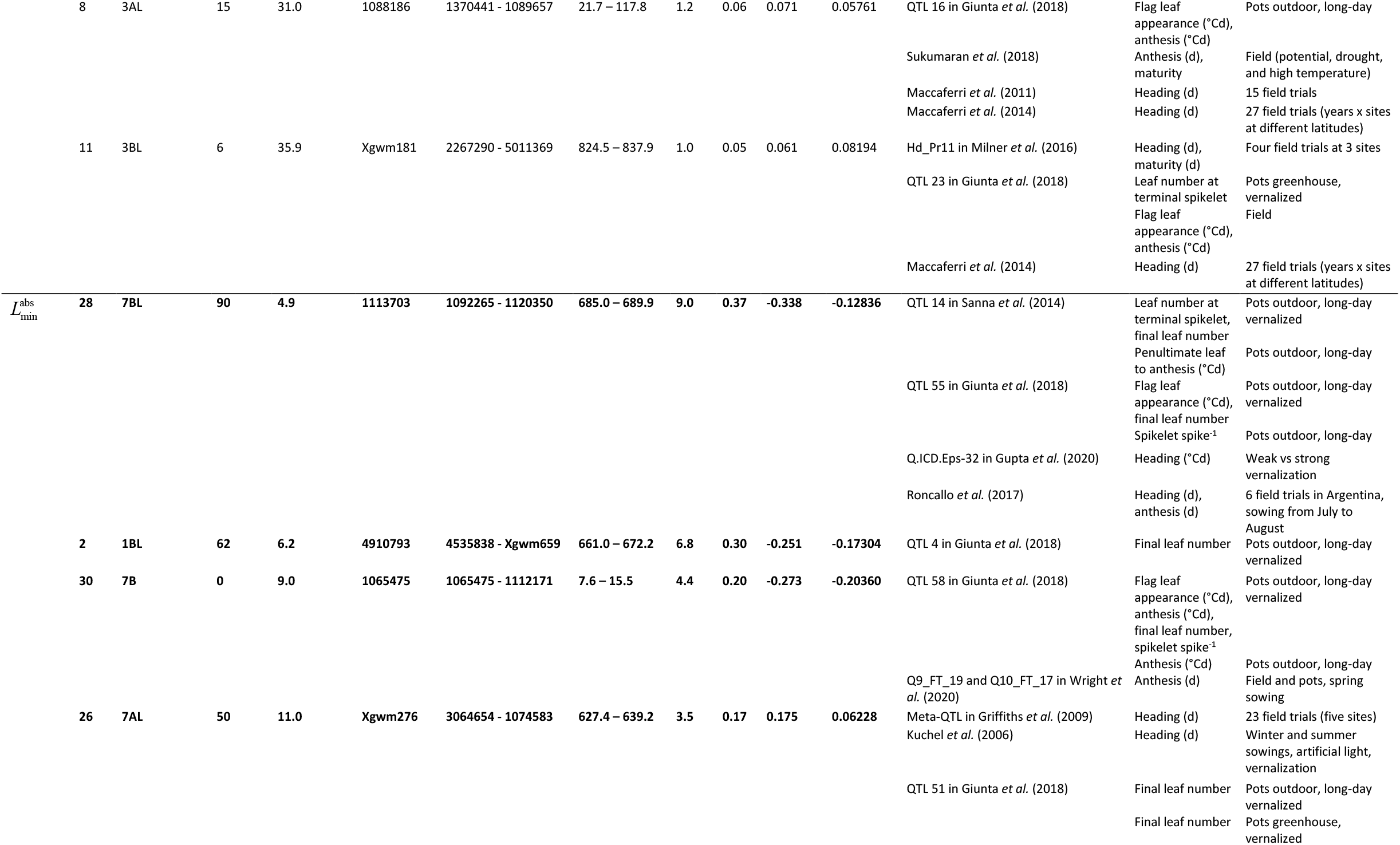
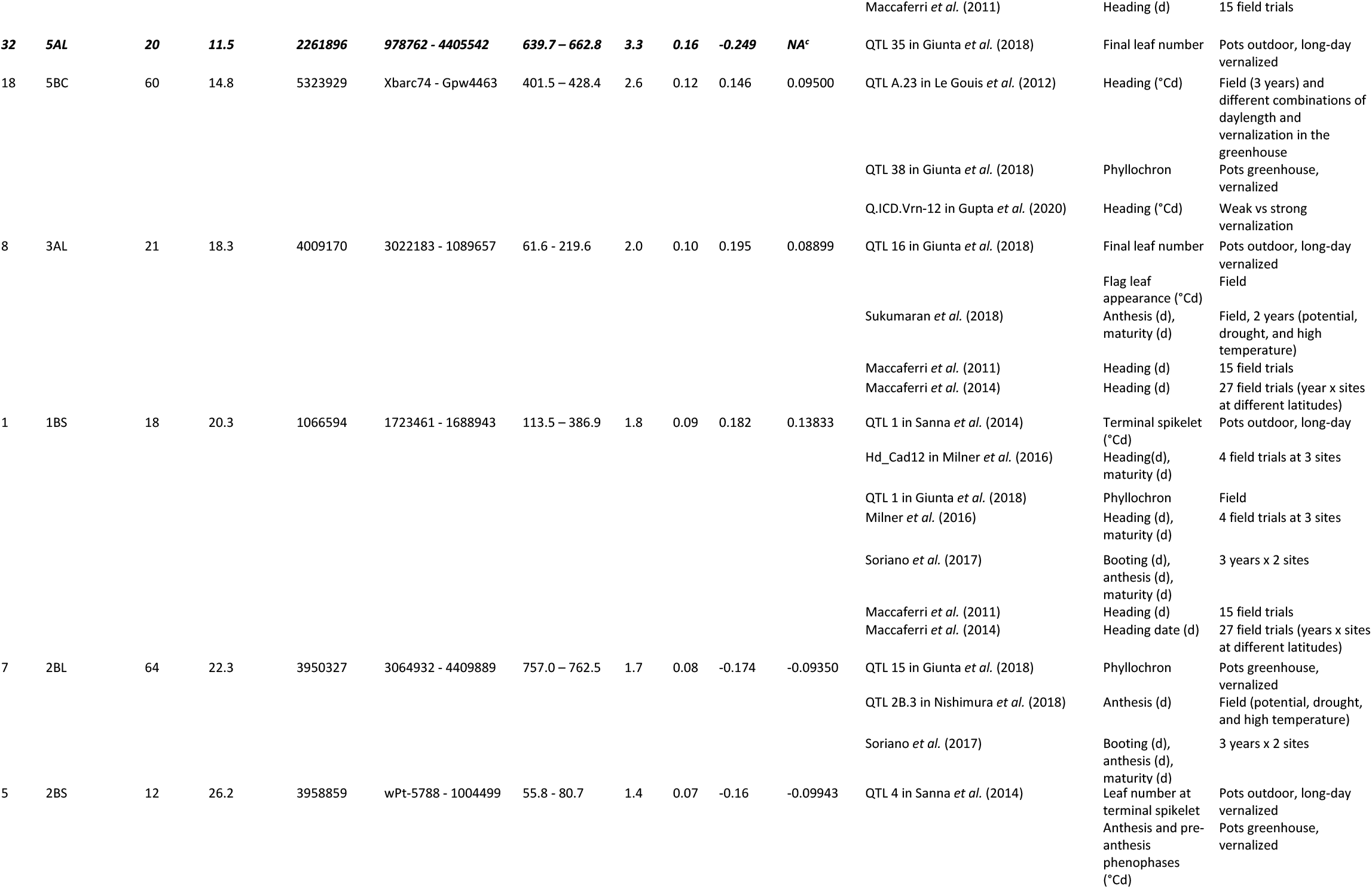
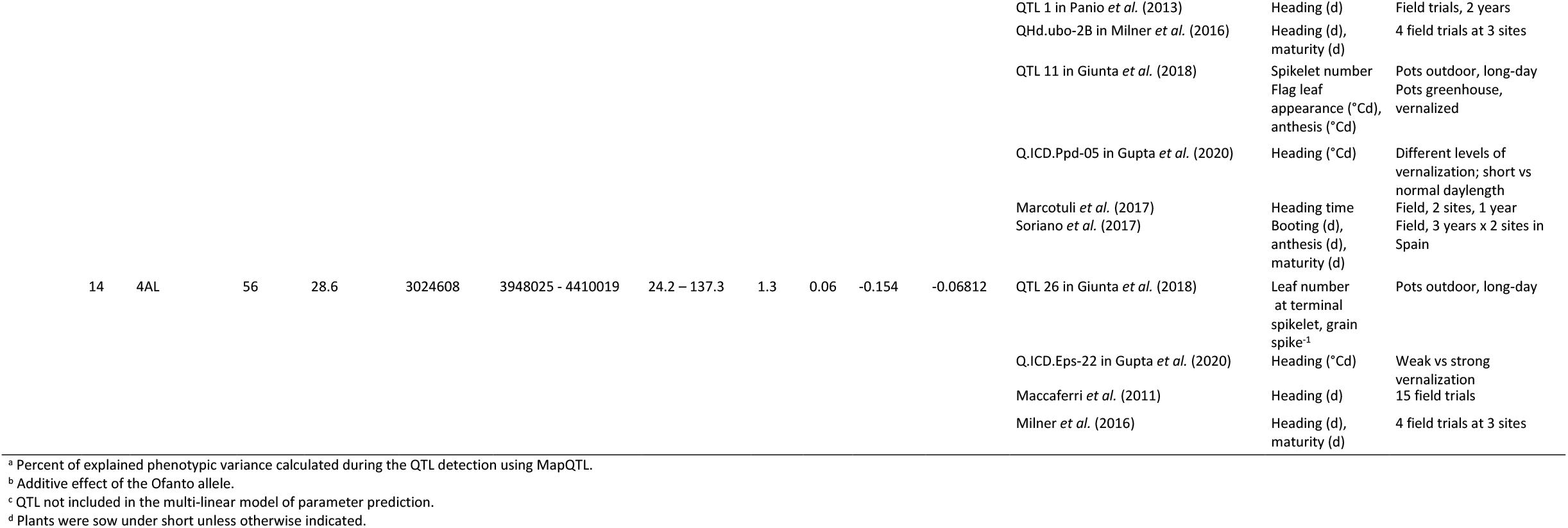
QTL used to predict the five genetic parameters of *SiriusQuality* phenology model. Two moderate QTL (QTL 31 and 32) not used to predict SLDL and 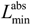 are also indicated in italic face. *P*, phyllochron; SLDL, daylength sensitivity; VAI, rresponse of vernalization rate to temperature; 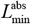, absolute final leaf number; PFLLAnth, Phyllochronic duration of the period between flag leaf ligule appearance and anthesis. Major (LOD < 5 and *r*^2^ > 0.1) and moderate (LOD > 2.8) QTL are indicated in bold face.

Two moderate QTL (LOD > 2.8) for 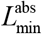 colocalized with known developmental genes (Fig. 3); QTL30 colocalized with Vrn-B3, and QTL32 with Vrn-A2 and FT-A5. *Vrn-A2* was also close to QTL16 for SLDL but not within the QTL confidence interval. We also found one tentative QTL for 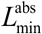 (and SLDL), QTL5, that colocalized with Ppd-B1 loci. For VAI, the major QTL15 colocalized with Vrn-A1 on chromosome 5A, and the peak marker for two tentative QTL, QTL1 and QTL8, colocalized with CO-B9 and FT-A2, respectively. The peak marker of QTL23 for *P* colocalized with CO-B2 locus. For the other two parameters, PFLLAnth and SLDL, the only associations to known developmental genes regarded putative QTLs. For PFLLAnth, QTL25 colocalized with Co-A1 locus and for SLDL, the peak marker of QTL2 and QTL5 colocalized with ELF-B1 and Ppd-B1 loci, respectively.

The five genetic parameters of *SiriusQuality* were estimated using the 79 QTL with a LOD score > 1. Eleven significant QTL and 21 tentative QTL with a LOD score value between 1 and 2.8 were used as predictors in the fitted statistical models (Table 2). *P*, SLDL, VAI, PFLLAnth and, 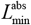 were predicted with 11, 10, 8, 6, and 10 QTL, respectively. QTL 32, which collocated at *Vrn-A2* was not selected in the multilinear model to predict 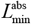, but the tentative QTL16, close to *Vrn-A2*, was used to predict SLDL. Seven tentative QTL collocated with several parameters. Tentative QTL8 and QTL14 were associated with four of the five parameters, the other five tentative QTL (QTL1, QTL5, QTL7, QTL10, and QTL25) were associated with two parameters.

The coefficients of the multi-linear model (Table 2) were well correlated with the additive effect of the QTL (all *r*^2^ > 0.87 and *P* < 0.002), except for SLDL (*r*^2^ = 0.01, P = 0.56). Thirty one of the 33 of the tentative QTL used to predict the parameters colocated with known QTL for heading date or other wheat phenology traits (Table 2). The fitted multi-linear model predicted the five parameters without significant bias (Fig. 3), they explained 36% (for PFLLAnth) to 63% (for *P* and 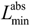) of the genotypic variation of the parameters. The relative RMSE for *P*, SLDL, VAI, PFLLAnth and, 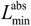 were 1.7%, 18.9%, 30.7%,9.6%, and 4.1%, respectively. The QTL-based parameters of the two parents of the RILs were also well estimated, especially for Cappelli (Fig. 3).

### Predictions of leaf stage

As illustrated in Figure 4 for the lines with the highest (135.9 leaf °Cd^-1^) and lowest (118.6 leaf °Cd^-1^) values of *P*, the model parametrized with the estimated (original) parameters predicted well the rate of main stem leaf appearance for the treatment LDV used to estimated *P* (Fig. 2A) but also for the treatments not used to estimate it (SDB, LDNV; Fig. 2C,E), as well as for the field experiment OT13 (Fig. 2G). For the latter experiment, the RMSE for main stem leaf number was only 0.15 leaves (Table 2). The QTL-based model also predicted well the rate of leaf appearance in all treatments (Fig. 2C, D, F, and H), and the RMSE for the validation experiment was close to that of the model with the original parameters (Table 2).

**Figure 2.**
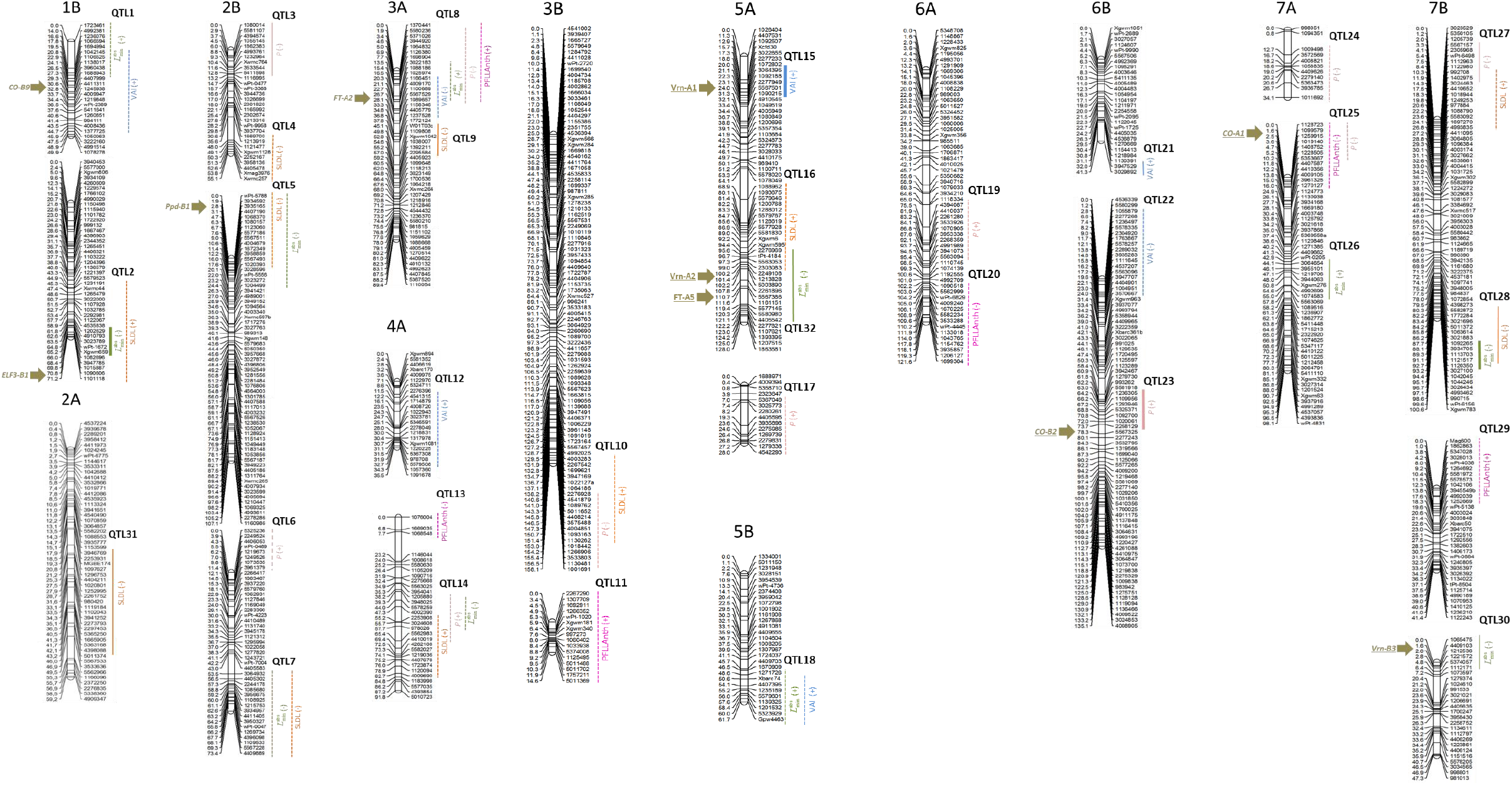
Chromosomal regions harboring QTL for the five genetic parameters of the *SiriusQuality* phenology model for the Ofanto × Cappelli RILs population. Genetic distances (cM) are indicated on the left of each linkage group, marker codes are indicated on the right. The vertical bars indicate the 95% confidence intervals (CI). Dashed CI bars indicate tentative QTL with 1 < LOD < 2.8; solid CI bars indicate moderate QTL with 2.8 < LOD < 4.9; thick solid CI bars indicate major QTL with LOD > 5. Signs in parenthesis after the parameter names indicate the sign of the additive effect of the Offanto allele. Major phenology genes in segregation in the population are indicated by horizontal arrows on the left of the linkage groups.

### Predictions of Final leaf number

The treatments in the calibration experiment had large effects on *L*_f_. As expected, on average *L*_f_ was the lowest for LDV (averaging 9.0 leaves) and the highest for LDVN (averaging 13.6 leaves; Fig. 3A). The genetic variability of *L*_f_ was also much higher for the LDNV-grown plants than for the two other treatments. The model explained 90% of the genotypic variation of *L*_f_ for the mean of the three treatments (Table 2) but only 35% for SDV. For the field experiment of the validation data set where *L*_f_ was recorded (OT13), the RMSE was only 0.46 leaves, but the model explained 20% of the genotypic variance. The RMSE for *L*_f_ was about two-times higher for the QTL-based model than for the model with the estimated parameters. The higher error of the QTL-based model was mainly due to a higher lack of correlation (Table 2). However, for validation data set both models gave similar results.

**Figure 3.**
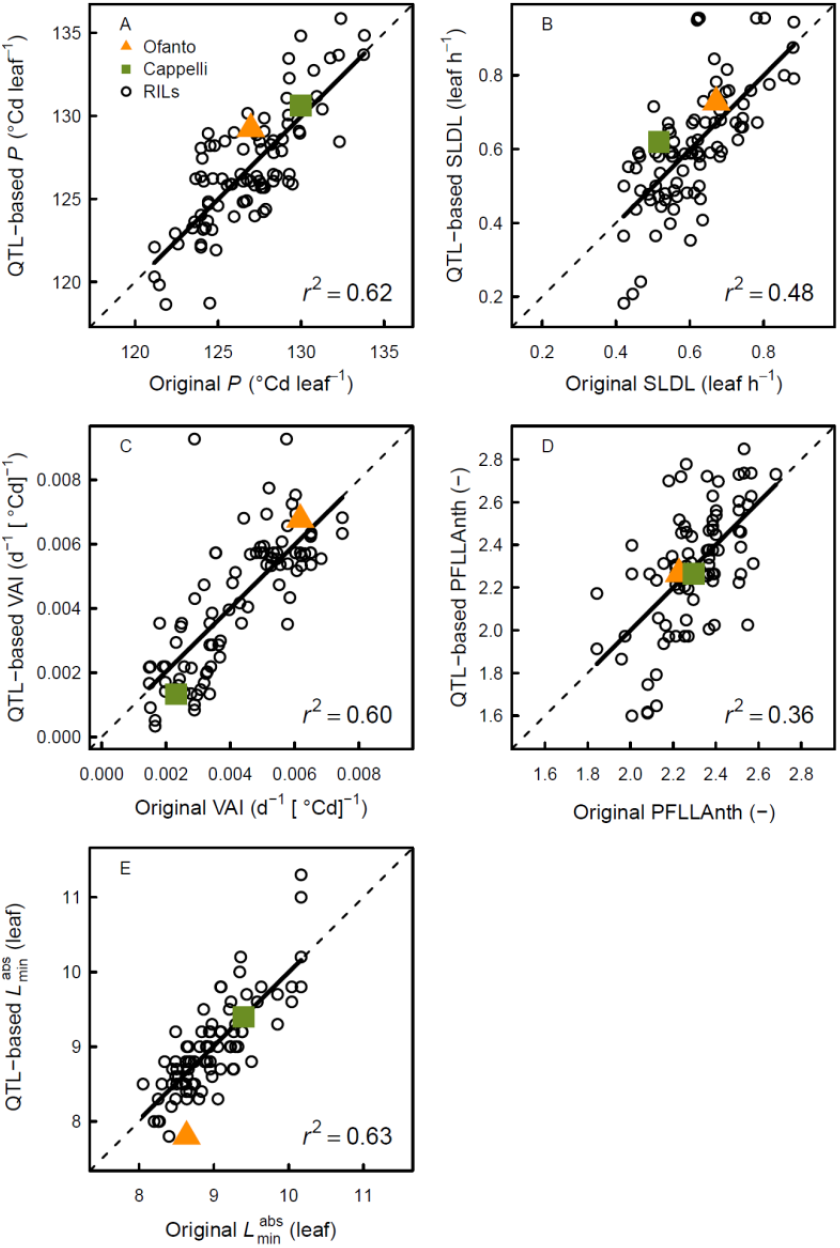
QTL-based versus original estimations of the five genetic parameters of the *SiriusQuality* phenology model for 91 RILs of the Ofanto (Of) × Cappelli (Ca) cross. The phyllochron (*P*), the sensitivity to day length (SLDL), the response of the vernalization rate to temperature (VAI), and the number of phyllochron between flag leaf ligule appearance and anthesis (PFLLAnth) were calibrated using the three environments of the calibration dataset, while the absolute minimum leaf number 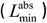 was measured in the LDV treatment. Dashed lines are 1:1 lines and solid lines are linear regressions. Note that the two parents were not used for QTL identification.

### Predictions of flag leaf ligule appearance date

In the calibration experiment, the average number of days between seed imbibition and the appearance of the flag leaf ligules was 56, 73, and 135 for the LDV, SDV, and LDNV, respectively (Fig. 3B). The shorter duration for LDV compared with LDNV was due to the low temperature during the vernalization treatment. The lower number of leaves for LDV compared to LDNV did not compensate for the low rate of leaf emergence during the vernalization treatment for LDNV. The model predicted the flag leaf ligule appearance date with a RMSE of 0.9 days for the mean of the three treatments used for parameter estimation (Fig. 3C, Table 2) and explained 60% (for LDV) to 99% (for SDV) of the genotypic variance. The RMSE was more than three-folds higher for LDNV and LDV than for SDV. For the validation trial for which the flag leaf ligule appearance was recorded (OT13), the RMSE was significantly higher (4 days) than for the calibration data set. The model explained only 28% of the genotypic variance for flag leaf ligule appearance date for OT13 (Table 2, Fig. 4C), which was mainly responsible for the model error (LC accounted for 70% of the MSE).

**Figure 4.**
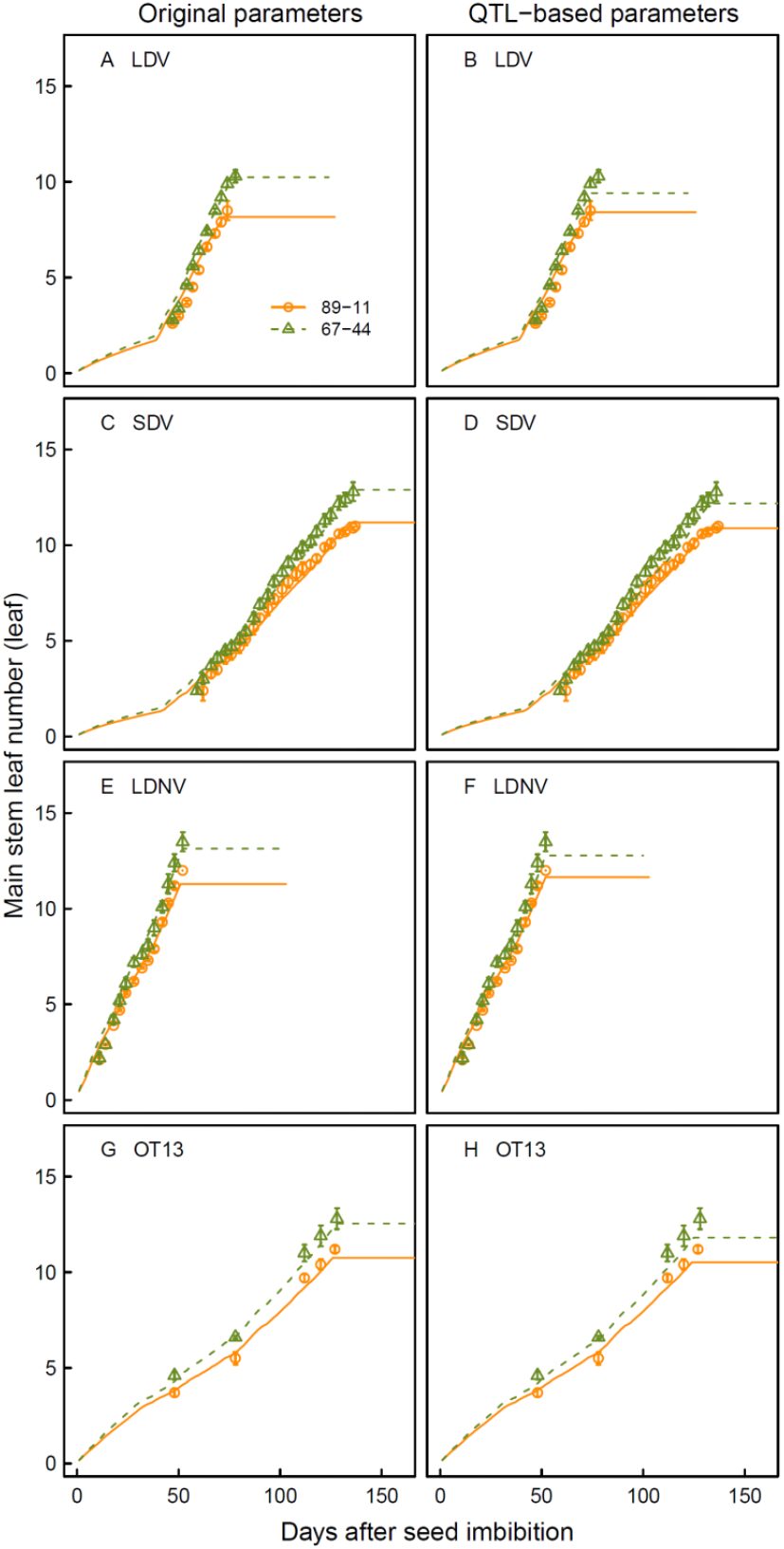
Haun stage versus days after anthesis for the two RILs of the Ofanto × Cappelli cross with the highest (89-11) and lowest (67-44) phyllochron for the calibration (A-F) and the validation (G-H) data sets. Symbols are measurements, lines are simulations. The names of the experiments as defined in Table 1 are given in the figure. Simulations were performed with the wheat model *SiriusQuality* using the original (A, C, E, and G) and QTL-based (B,D, F, and H) genetic parameters. Measurements are the mean ± 1 s.d. for *n* = 4 independent replicates.

For LDV, the RMSE for the days to flag leaf ligule appearance were similar for the QTL-based model and the model with the estimated parameters, while for the LDNV and SDV it was about two- and five-times higher for the QTL-based model than for the model with the estimated parameters. For the validation data (OT13), the RMSE of both models were similar, but the QTL-based model explained only 11% of the genetic variation of the date of flag leaf ligule appearance, compared with 65% for the model with the estimated parameters. The ranking of the lines was better conserved (ρ = 0.58 and 0.36 with the original and QTL-based parameters, respectively).

**Table 3.**
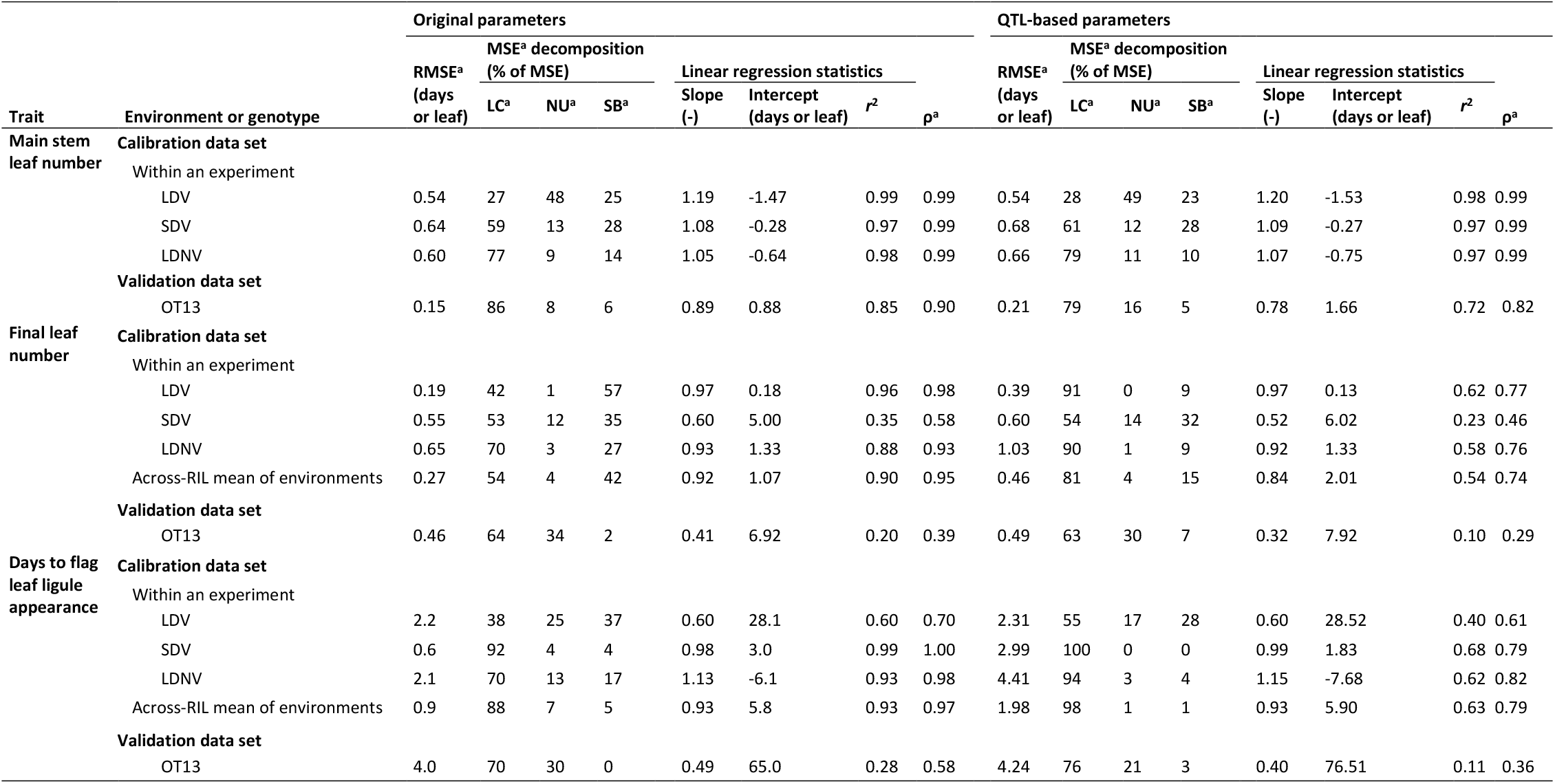
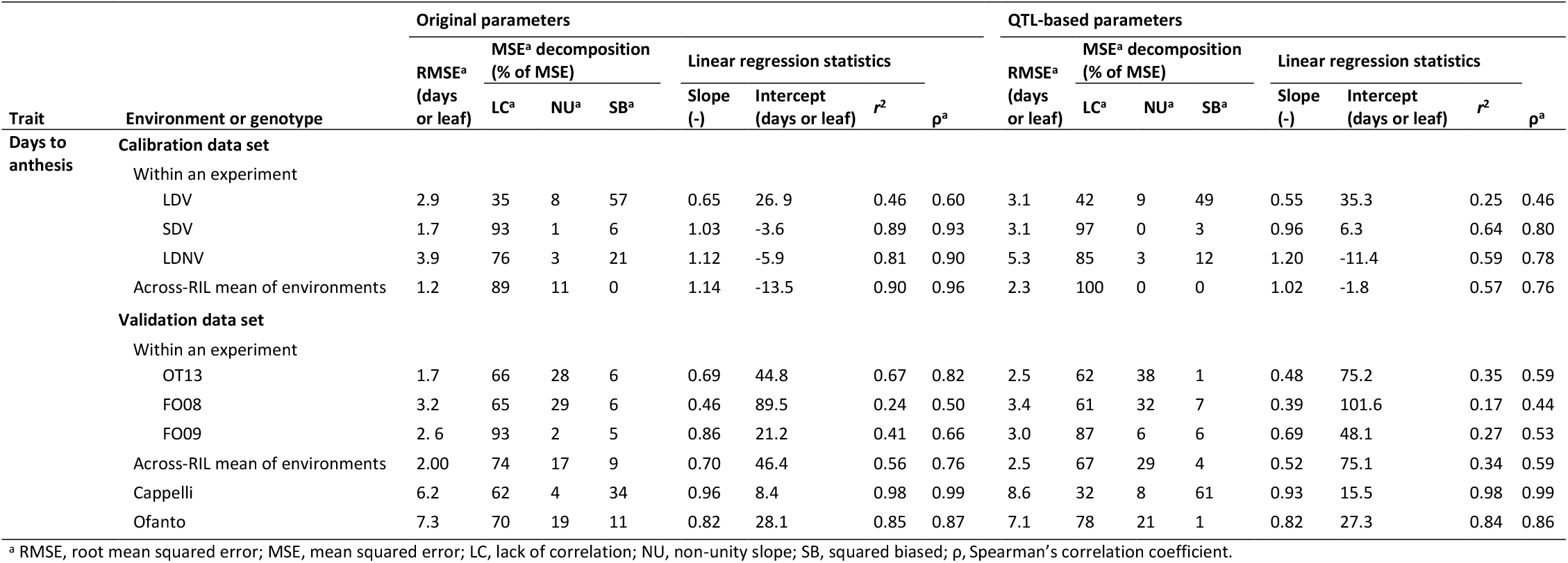
Statistics of model performance to predict days to flag leaf ligule appearance, final leaf number and days to anthesis using the original and the QTL-based parameters for the calibration and the validation data sets. Days to flag leaf ligule and anthesis were calculated from the day after seed imbibition and sowing for the calibration data set and validation data sets, respectively.

### Predictions of anthesis date

In the calibration experiment, the number of days to anthesis was about two-times higher for SDV than for the long day treatments (Fig. 3E). The genotypic variability was also much higher for SDV-grown plants. Although three of the five genetic parameters were estimated with the LDV treatment, the model explained less of the genotypic variance for this treatment than for the other two (Table 2).

Across the three treatments of the calibration experiment, the RMSE for anthesis date ranged from 1.7 (SDV) to 3.9 (LDNV) days and the *r*^2^ ranged from 0.46 (LDV) to 0.89 (SDV). In the three independent field experiment, the RMSE and *r*^2^ for the mean anthesis across the RILs were 2.0 days and 0.56, respectively. In OT13 and FO08, the model error was mainly due to a lack of correlation, while in FO09 about half was due to a lack of correlation and non-unity slope.

For the validation data set, the RMSE for anthesis date was 0.2 to 0.8 days higher for the QTL-based model compared with the model with the estimated parameters (Table 2, Fig. 6F). On average over the three experiments of the validation data set, the QTL-based model explained 34% of the genetic variation of anthesis date, which is slightly more than half of the genetic variation explained by the model with the estimated parameters. The ranking of the lines was more conserved between the estimated and QTL-based parameters (0.76 vs. 0.59).

**Figure 5.**
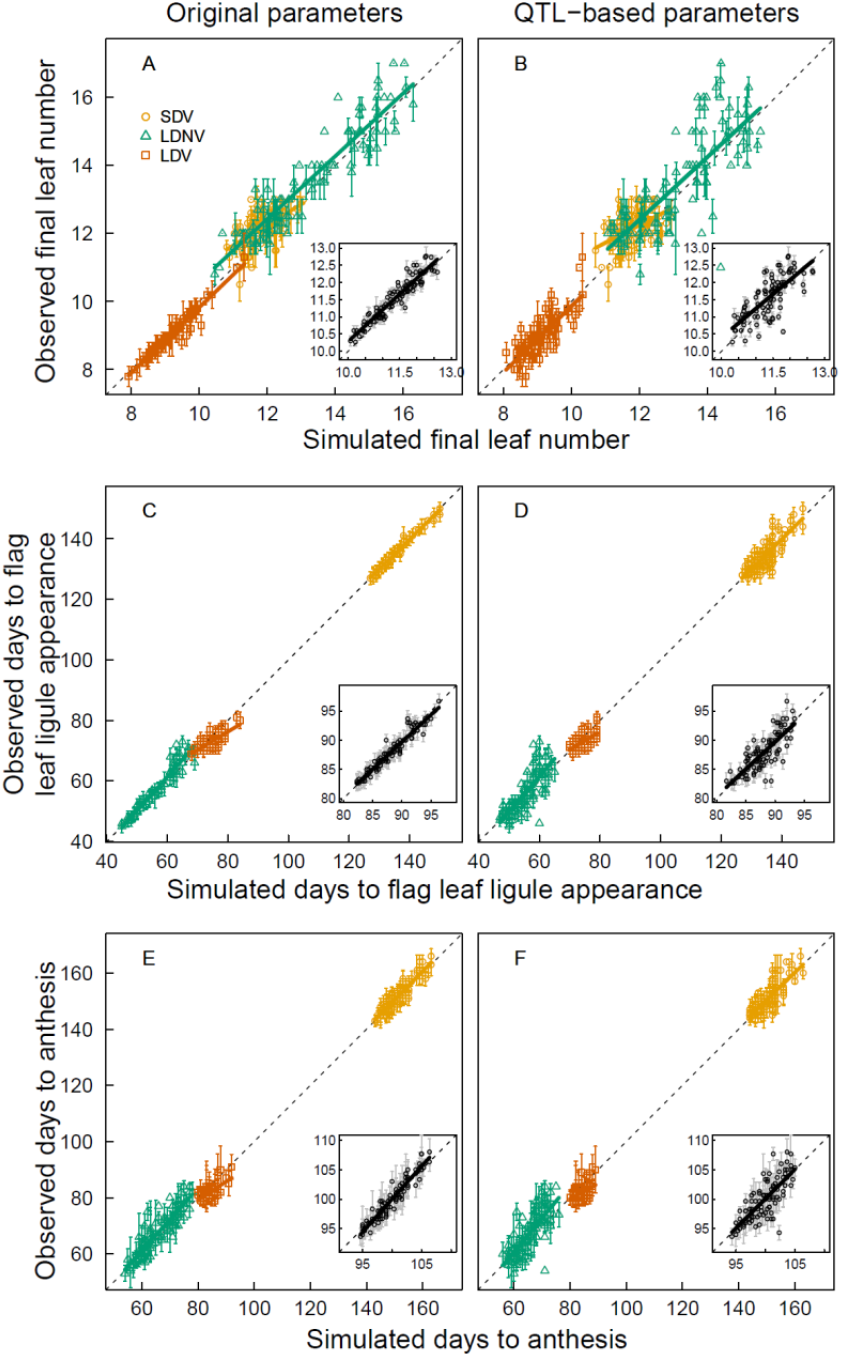
Observed versus simulated final leaf number (A and B), days to flag leaf ligule appearance (C and D) and days to anthesis (E and F) for 91 RILs of the Ofanto × Cappelli cross. Data are for the short days vernalized (SDV, circles), long days non vernalized (LDN, triangles), and long days vernalized (LDV,squares) treatments of the experiment used to estimate the genetic parameters of the *SiriusQuality* wheat phenology model. Simulations were performed using original (A, C, and E) and QTL-based (B, D, and F) genetic parameters. Inset panels show the mean values for the three experimental treatments. Days to flag leaf ligule and anthesis were calculated from the day after seed imbibition. Dashed lines are 1:1 lines, solid lines are linear regression. Measurements are the mean ± 1 s.d. for *n* = 4 independent replicates.

**Figure 6.**
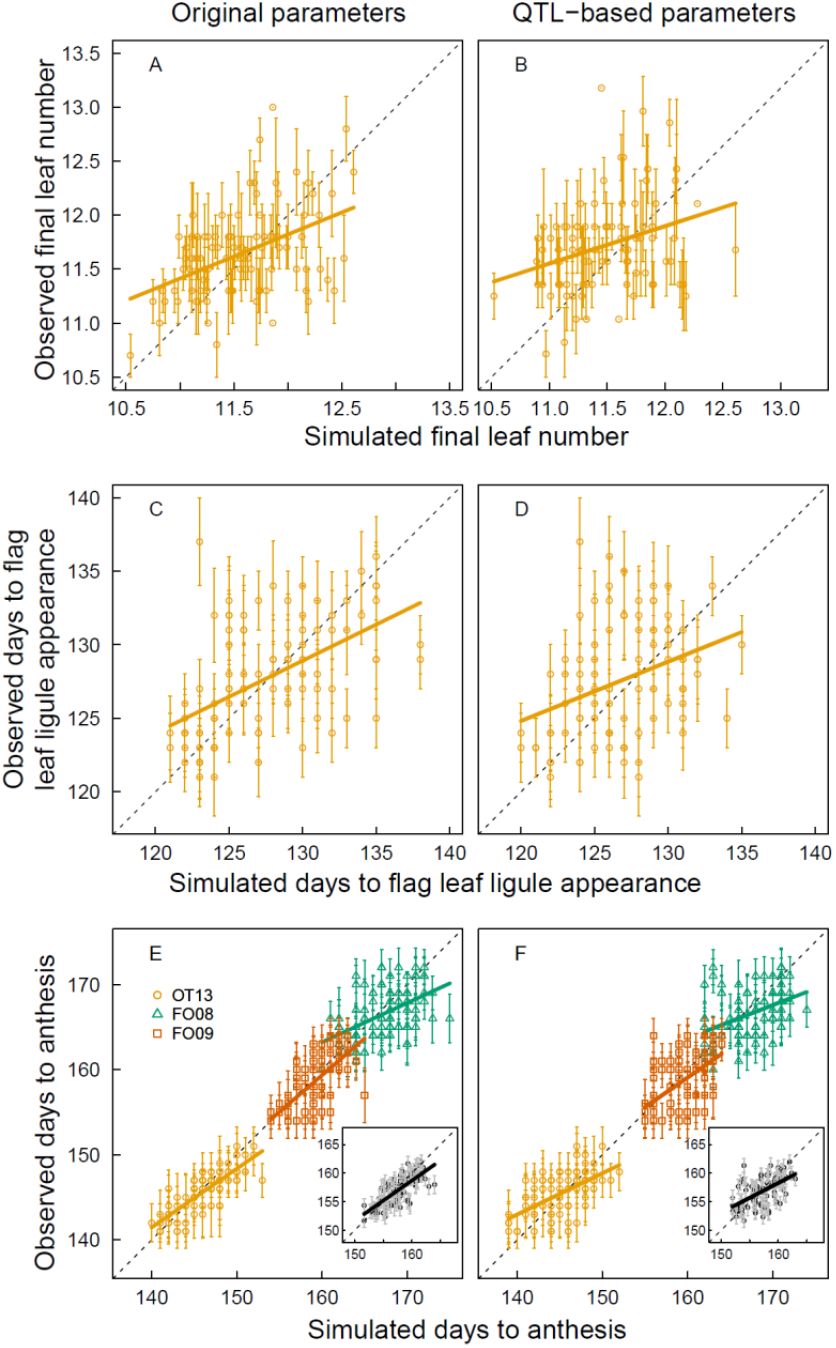
Observed versus simulated final leaf number (A and B), days to flag leaf ligule appearance (C and D) and days to anthesis (E and F) for 91 RILs of the Ofanto × Cappelli cross grown in the field in Ottava, Sardinia, Italy during the 2021-2013 growing season (OT2013, circles) and in Foggia, Italy during the 2007-2008 (FO08, triangles) and 2008-2009 (FO09, squares) growing seasons (validation data set). Simulations were performed with the SiriusQuality wheat phenology model using original (A) and QTL-based (B) genetic parameters. Final leaf number and days to flag leaf ligule appearance were recorded in OT2013 only. Inset panels in (E) and (F) show the mean values for the three field experiments. Days to flag leaf ligule and anthesis were calculated from the day after sowing. Dashed lines are 1:1 lines, solid lines are linear regression. Measurements are the mean ± 1 s.d. for *n* = 3 independent replicates.

### Predictions of anthesis date for new genotypes in new environments

The QTL-based model was further evaluated for the two parents of the RIL population grown in the field in experiments not used for parameter estimation. The two parents were not used to identify QTL, it is thus a test of the ability of the QTL-based model to predict new genotypes. Across all site/year/sowing date combinations, the number of days to anthesis ranged from 71 to 170 days for Cappelli and from 94 to 171 days for Ofanto. The model with the original parameters predicted anthesis date for Cappelli and Ofanto with a RMSE of 6.2 and 7.3 days and a *r*^2^ of 0.98 and 0.85, respectively (Table 2, Fig. 7A). The RMSE of the QTL-based was higher than that of the original model by 2.4 days for Cappelli and was similar for both models for Ofanto (Table 2, Fig. 7B). For Cappelli, the model with both the original and QTL-based parameters had a larger RMSE for the autumn sowing dates (late November – mid December) than for the spring sowing dates (late January – late March). For the QTL based model, the RMSE and *r*^2^ were 7.7 d and 0.97 for the autumn sowing dates and were 9.9 days and 0.59 for the spring sowing dates, respectively.

**Figure 7.**
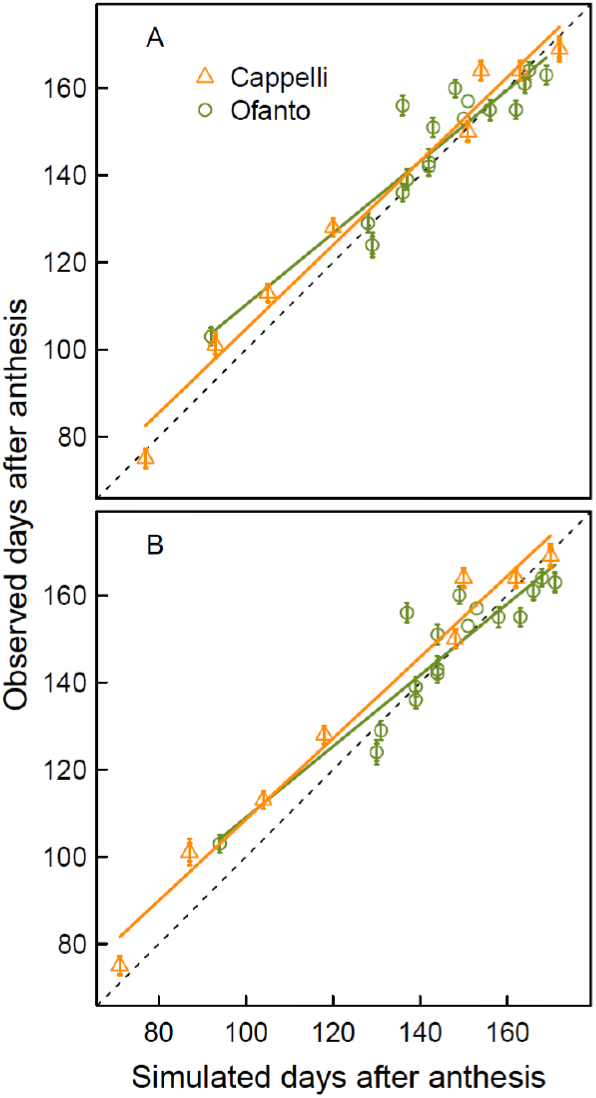
Simulated versus observed days to anthesis for the two parents grown in the field in 18 (Cappelli) and eight (Ofanto) site/year/sowing date combinations. Simulations were performed with the wheat model *SiriusQuality* using the original (A) and QTL-based (B) parameters. Days to flag leaf ligule and anthesis were calculated from the day after sowing. Dashed lines are 1:1 lines, solids lines are linear regression. Measurements are the mean ± 1 s.d. for *n* = 3 independent replicates.

## Discussion

Gene- or QTL-based models are useful to integrate ecophysiological, genetic and molecular knowledge and to improve simulation models. They are also powerful tools to predict genotype performance (Chenu *et al*., 2009), identify ideotypes (Bogard *et al*., 2020b) or combinations of alleles or loci (Bogard *et al*., 2020a; Zheng *et al*., 2016) to adapt genotypes to target environments under current or future climate scenarios, or to design new crop management strategies for specific existing or virtual (new combinations alleles or loci associated with model parameters) genotypes (Martre *et al*., 2014). In this study, we used a model that integrates our current understanding of the physiology of wheat development and phenology to predict the development and phenology of a RILs population of durum wheat with parameters estimated with vernalization and photoperiod treatments. We identified major or moderate QTL associated with four of the five genotypic parameters of the model. We then used this genetic information to estimate the value of parameters and to predict plant development and anthesis date of the RIL population, including the parents, which were not used for QTL identification, in new environments in the field. We discuss the approach we used to estimate the parameters of the model and their association with QTL and major phenology genes that collocate at QTL.

### Genotypic parameters for earliness *per se*, cold requirement, and photoperiod sensitivity can be estimated independently with vernalization and photoperiod treatments

We estimated five genotypic parameters independently for earliness *per se*, cold requirement, and photoperiod sensitivity using three vernalization and photoperiod treatments. This procedure minimized the risk of finding local minima and reduced the computation time for parameter estimation. It increases the risk of compensation for errors, but it is a better test of the model compared with the estimation of all parameters together.

For the validation data set, the RMSE for anthesis date was low and was similar for the model with estimated parameters (2.0 d RMSE) and with QTL-based parameters (2.5 d RMSE). Compared with previous studies, the RMSE for anthesis date, was lower than that reported for wheat (5 to 8.6 d in Bogard *et al*., 2014; 6 to 9 d in White *et al*., 2008; 4.3 d in Zheng *et al*., 2013) or other species (5 to 7.5 d in Messina *et al*., 2006 for soybean; 7.6 to 15 d in Uptmoor *et al*., 2012 for Brassica oleracea; 4.2 d in Uptmoor *et al*., 2017 for spring barley). As in all these studies, we found a significant decrease of the percentage of genetic variations explained with the QTL-based parameters (34%) compared with the estimated original parameters (56%). The ranking of the lines for the time to anthesis was better conserved than the *r*^2^, the Spearman’s rank correlation coefficient was 0.76 with the estimated parameters and 0.59 with the QTL-based parameters. The lower performance of gene- or QTL-based models can be due to undetected effects of minor QTL (Yin *et al*., 2005), poor estimation of allelic effects of known QTL (Uptmoor *et al*., 2012), the use of markers outside the causal polymorphism and possible recombination between markers in linkage disequilibrium (Bogard *et al*., 2014), or the method used to estimate the QTL or gene parameters (Zheng *et al*., 2013), in addition to the errors of the model itself.

Bogard *et al*. (2014), calibrated an empirical phenology model modified from Weir *et al*. (1984) for a panel of 210 bread wheat genotypes. They estimated the parameters of their model using heading date data from field trials sown in the autumn and spring for the winter and spring type genotypes, respectively. For the winter type genotypes, they found several combinations of parameters that gave similar simulation results for anthesis date and the overall (for spring and winter types) RMSE for heading date was on average two-folds higher for the spring than for the autumn sowings. He *et al*. (2012) calibrated the model used here for 16 winter wheat cultivars with field data form autumn sown crops and concluded that VAI cannot be estimated using only autumn-sown field trials, even with a large number of environments with a wide range of winter temperature and latitude. These studies clearly indicate that to estimate vernalization parameters, vernalization and daylength treatments are needed, either in the field or under controlled conditions, as used in this study and in previous studies (Yin *et al*., 2005; Zheng *et al*., 2013). Here we show that a minimum of three treatments is required to estimate the three components of phenology.

The treatments should allow for a complete satisfaction of cold requirement of all the studied genotypes. In our study, in the long day vernalized treatments *Lf* varied between 7.8 and 11.3 leaves among the lines, while the minimum number of leaves of vernalized spring wheat genotypes is around 6 leaves (Levy and Peterson, 1972). 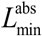 was thus likely overestimated because at least some lines were not fully vernalized in the SDV treatment. This may explain the negative correlation we found between 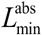 and SLDL and the five common non-significant QTL for these two parameters. This hypothesis is also supported by the colocation of QTL32 for 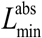 at *Vrn-A2. VRN2* is a floral repressor expressed only under long days, where it delays flowering until plants are vernalized by repressing *VRN3* (Trevaskis *et al*., 2007). During cold periods the induction of *VRN1* represses *VRN2*, allowing the day-length response (Yan *et al*., 2004). Therefore, the colocation of QTL32 for 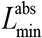 at *Vrn-A2* can be explained by admitting that the vernalization treatment in the SDV treatments resulted in some lines being not fully vernalized.

We used twice-weekly measurements of LS, final main stem leaf number, and the date of anthesis of long-day vernalized plants to estimate the three earliness *per se* parameters (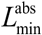, *P*, and PFLLAnth), while the rate of vernalization (VAI) and the sensitivity to daylength (SLDLL) were estimated using observations of the date of flag leaf ligule appearance of nonnvernalized plants grown under long days (LDNV) and vernalized plants grown under short days (SDV), respectively. SLDL and VAI were estimated by minimizing the error for the date of flag leaf ligule appearance rather than for *L*_f_ to reduce the compensation for error for PFLLAnth. It also improved the simulation of the stage flag leaf ligule just visible, which is synchronous with the stage male meiosis, a key stage to model the impact of abiotic stress on grain number abortion (Barber *et al*., 2015).

Depending on the objectives of the study, our phenotyping protocol can be greatly simplified. For instance, the minimum information required to calibrate the model for spring genotypes are LS measured every about three leaves between leaf 3 and 9 and anthesis date for short- and long-day grown plants (Jamieson and Munro, 2000). To calibrate the model for winter wheat, the date of flag leaf ligule appearance or anthesis of nonnvernalized plants grown with long days is also required. With the rapid development of plant phenomics, all the measurements required to calibrate the model for new genotypes can be automatized at high throughput. High-resolution RGB imagery with deep-learning techniques has recently been used to estimate heading date (Madec *et al*., 2019), and, combined with three-dimension plant architecture models, LS, and thus *P*, can also be accurately estimated (Liu *et al*., 2019). It should also be possible to develop high-throughput phenotyping methods for the dates of flag leaf ligule appearance and anthesis using similar techniques. These methods would greatly facilitate the calibration of the model for large genetic panels for genetic analyses.

### Model parameters are to a large extent genetically independent and are associated with major phenology genes

We predicted the parameter values considering only the additive effect of the QTL but Bogard *et al*. (2014) found non-significant or small bi-locus marker x marker interactions for markers associated with model parameters for vernalization requirement and photoperiod in the bread wheat panel they studied. Our objective was not to identify robust QTL but to predict the genetic value of parameters; therefore, we used all available information and predicted the parameters using all (tentative) QTL with a LOD score > 1.

The multi-linear models predicted the five genotypic parameters with six to 11 QTL and explained 36% to 68% of the genetic variation of the estimated parameters. In comparison, Bogard *et al*. (2014) estimated three model parameters and their multi-linear predictions based markers explained 68% to 71% of the variation of their parameter. Recombinations between markers may be the cause of the large part of the genetic variation of the parameter not explained by the QTL in our study. The remaining unexplained variations of the parameters may be due to QTL with smaller effect that were not detected because of the limited size of our population and insufficient coverage of the genetic map.

Twenty-nine of the 30 QTL used to predict the parameters colocalized with known phenology QTL. Our study provides a quantification of their effect that is independent of the environment that can be used to predict the phenology of genotypes in different environments. They also provide new insights onto the physiological processes controlled by the associated regions. Twelve of the 13 major and moderate QTL we identified were associated with only one parameter and several collocated at major phenology (Vrn-A1, Vrn-A2, Vrn-B3, Pppd-B1, CO-2, and FT-A5), reflecting that the parameters are genetically independent for the most part and that the model discriminates well the effect of the physiological processes controlling the phenological development of wheat.

PFFLAnth had a relatively high standard deviation between lines (0.28 phyllochron) but a low heritability (8.7%) and we found no significant QTL for this parameter. A previous study on the same population also did not find any significant QTL for the duration in thermal time between flag leaf ligule appearance and anthesis (Sanna *et al*., 2014). It has been reported that this period is sensitive to daylength (Fischer, 2011; Whitechurch *et al*., 2007). Here, PFFLAnth was significantly correlated with *P* and SLDL (Fig. 1). These correlations were, at least in part, due to the nature of these parameters and the way they were estimated. *P* and SLDL directly depend on the rate of leaf appearance, and PFLLAnth is expressed in phyllochronic time. The impact of a different rate of leaf appearance induced by daylength is mediated by the number of plastochrons that the plant is able to produce and by the variation in duration induced by photoperiod. Improving the prediction of the duration of the phase between flag leaf appearance and anthesis (that is PFFLAnth) is an important model improvement target as it has a strong effect on grain number per ear (Fischer, 2011).

In contrast with major and moderate QTL, half of the tentative QTL were associated with two to four parameters (Fig. 8). Four of these QTL, and the tentative QTL28, were associated with 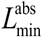 and SLDL. At least some of these QTL colocations are likely related to incomplete vernalization of some lines in LDV treatment (e.g. the common QTL between PFLLANTh and VAI). SLDL and *P* were significantly correlated (*r* = 0.40, *P* = 0.001) and we found two tentative QTL (QTL10 and QTL27) associated with these two parameters (Figs. 3, 8). In winter barley, under long days conditions genotypes carrying the photoperiod sensitive alleles of *Ppd1-H1* (early flowering) have a reduced leaf length and an higher leaf appearance rate (Digel *et al*., 2016). In wheat, the daylength insensitivity alleles of *Ppd-1* was also found to reduce phyllochron under long day in the field but only after leaf 7 (Ochagavía *et al*., 2017), confirming the effect of photoperiod on the rate of emergence of late-formed leaves found by Miralles and Richards (2000)In agreement with these results, QTL10 and QTL27 had opposite additive effects on SLDL and *P*. These results suggest the opportunity to consider an effect of daylength sensitivity on *P*. Although expressed only for the last leaves, this would modify the duration of the terminal spikelet to anthesis and flag leaf appearance to anthesis periods. Although none of the mentioned QTL collocated at *Ppd-1*, they may carry genes down- or up-stream of *Ppd-1*. However, the common genetic determinism of *P* and SLDL need to be further studied as we cannot rule out that it can be driven by carbon limitations during the stem extension period (Baumont *et al*., 2019).

**Figure 8.**
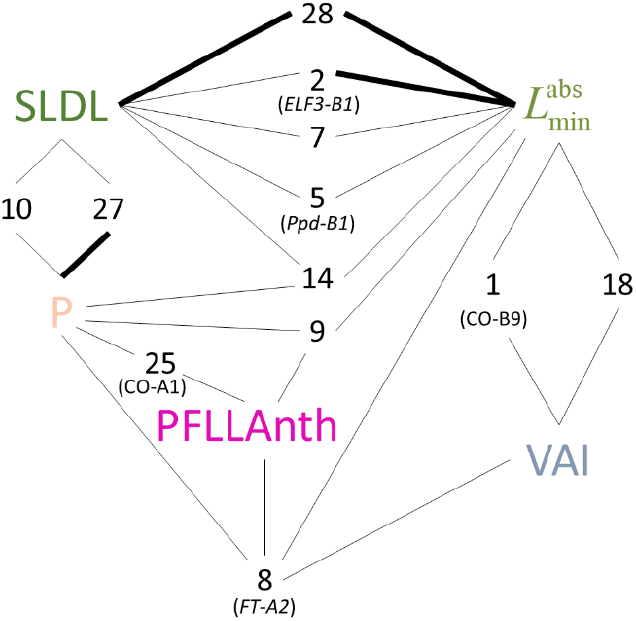
Schema of the QTL associated with two or more model parameters. Tick lines are major and moderate QTL with LOD > 2.8 and thin lines are tentative QTL with LOD between 1.0. and 2.8. Numbers correspond to the QTL numbers in Table 2 and in Figure 3. Parameters are defined in Table 1. Major phenology genes that collocate at QTL are indicated under the QTL numbers.

In conclusion, The QTL-based model of phenology developed in this study gives the possibility to quantify the effect of major phenology genes on agronomically important traits that are to a large part determined by phenology (e.g. cold hardness, tillering, leaf size, plant height, and grain number per ear; Hyles *et al*., 2020) in diverse environments. In contrast with empirical models that simulate thermal times between phenological states, the model used in this study simulates key developmental stages (floral initiation, terminal spikelets, flag leaf tip and ligule appearance) that define phase switch changes in leaf area (Martre and Dambreville, 2018), tillering (Abichou *et al*., 2018), and spikelet production and floret abortion (González *et al*., 2011). Future model development should consider the rate and duration of the phases of spikelet primordium formation and floret development, which are controlled by flowering time regulators (Gol *et al*., 2017), and determine the number spikelet per ear and floret survival and abortion (González *et al*., 2011). Kirby (1990) showed that the rate of spikelet primordium formation is directly related to *L*_f_. In this study, we identified four major QTL for three parameters (*P*, SLDL, and 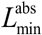) that colocalized with known QTL for spikelet number per ear (Table 2). Future studies with the model used in this study should also try to use makers in the causal polymorphism of known major phenology genes. This will provide quantitative information on the effect of this genes on important physiological traits (model parameters).

## Supplementary data

**Table S1.** Summary of the experiments used in this study.

**Table S2.** List of the species parameters of the wheat phenology model SiriusQuality used in this study.

## Acknowledgement

PM acknowledges the support of the University of Sassari during his stays to conduct this research through its 2020 visiting Professor program funded by the Regional Government of Sardinia, and Dr. Renaud Rincent (UMR GQE, INRAE, France) for helpful discussions.

## Author contributions

PM, RM, and FG designed the research, RM and FG designed and conducted the experiments; PD conducted the validation experiments at Foggia; AMM conducted the QTL analysis; AMM, PD, and DM mapped major genes on the genetic map and comparison QTL with known QTL; PM estimated the parameters, did the simulations, analyzed the data, and wrote the manuscript; all authors contributed to the revision of the manuscript.

## Data availability

The source code and the binaries of *SiriusQuality* can be freely downloaded at https://forgemia.inra.fr/siriusquality. The source code and the binaries BioMA component of the *SiriusQuality* phenology are available at https://doi.org/10.5281/zenodo.2478791.

